# The petal identity gene PhDEF has a major binding and regulatory action in the epidermis

**DOI:** 10.64898/2025.12.10.691030

**Authors:** Emma Désert, Quentin Cavallini-Speisser, Evelyne Duvernois-Berthet, Pierre Chambrier, Patrice Morel, Brice Letcher, Carine Rey, Jérémy Just, Suzanne Rodrigues Bento, Daniel Bouyer, Marie Monniaux

**Author notes:** These authors contributed equally (Emma Désert, Quentin Cavallini-Speisser).

## Abstract

In flowering plants, floral organ identity is specified by the combinatorial action of homeotic genes. While the role of these genes in the early specification of organ identity is well established, their late function throughout floral organ development and in specific cell types is much less characterized. In particular, since plant organs are structured in clonally-independent cell layers, whether and how homeotic identity interacts with cell layer identity is unknown. We have previously identified cell layer-specific mutants for the petal identity gene *PhDEF* in petunia flowers, resulting in drastically different petal phenotypes whether *PhDEF* is expressed in the petal epidermis or in the mesophyll. In this study, using a combination of single-cell RNA-Seq and chromatin immunoprecipitation on *phdef* cell layer-specific mutants, we find that PhDEF regulates a different set of target genes in the petal epidermis and mesophyll, with a major regulatory action in the epidermis. We uncover a high diversity of binding profiles in PhDEF target genes, with a complex combination of layer-specific or non-specific binding sites, and a much more prominent binding of PhDEF in the epidermis than in the mesophyll. Our study highlights that floral homeotic genes like *PhDEF* can have different regulatory actions in different cell contexts, here different cell layers, demonstrating that cell layer identity indeed influences the regulatory processes underlying homeotic identity.

## INTRODUCTION

In flowering plants, homeotic genes define the identity of floral organs in a combinatorial manner, as outlined in the ABC model^1–3^. These genes encode master transcription factors, most of them from the MADS-box family, that regulate the expression of a wide set of genes throughout floral organ development until organs are mature. Additionally, floral organs are structured in different cell layers that are inherited from the three clonally-distinct layers (L1, L2 and L3) of the shoot apical meristem^4–7^. The action of homeotic genes is therefore superimposed on a pre-established cell layer identity, raising the question whether homeotic function is dependent or influenced by cell layer identity.

Petals are often key players in pollinator attraction, and in the case of our focus genus *Petunia*, this is typified by the bright pigmentation or strong scent emission from the conical cells of the limb epidermis, while the long tube observed in some petunia species limits nectar access to adapted pollinators^8,9^. In the model organism *Petunia x hybrida* (petunia), petal identity is defined by B-class MADS-box regulators only: PhDEF (*Petunia x hybrida* DEFICIENS), PhGLO1 (GLOBOSA1) and PhGLO2^10–12^. The encoded proteins form PhDEF/PhGLO1 and PhDEF/PhGLO2 obligate heterodimers that translocate to the nucleus, bind DNA and regulate gene transcription, as was reported for their homologs in *Antirrhinum* and *Arabidopsis*^13–15^. These complexes activate the expression of the *PhDEF*, *PhGLO1* and *PhGLO2* genes themselves, resulting in an auto-activation loop^13,14,11^. This loop, that is conserved in all eudicots, buffers stochastic noise, leading to a sharp transition to petal identity, and ensures the maintenance of high expression levels of these genes until petals are mature^16,17^. Other proteins, in particular the E-class SEPALLATA MADS-box proteins^18,19^, are part of higher-order complexes with the B-class heterodimers to define petal identity^20–22^. B-class genes, after defining petal identity, also participate in their development and in the differentiation of their cell types. Indeed, experiments of temporal inactivation of B-class genes in *A. thaliana* have revealed that, at intermediate stages of flower development, petals easily revert to a sepal-like identity^23^, suggesting that expression of B-class genes needs to be maintained over the course of petal development for normal petal identity specification. Similarly, inhibition of B- class gene expression at later stages of *A. majus* flower development causes a reduction in petal size, conical cell size and scent emission^24^, and a reduction in scent production and emission was also recently reported in petunia petals when *PhDEF* expression is repressed at anthesis^25^. This suggests that petal identity remains partly uncommitted until intermediate-to-late stages of flower development, and that B-class genes act to maintain this late specification of identity through the control of specific cell fates and traits.

In this study, we investigate the late role of the petal identity regulator PhDEF in the two cell layers of the petunia petal, stemming from our previous identification of cell layer-specific *phdef* mutants^26^. In petunia, petals derive from the L1 and L2 meristematic layers that will form the petal epidermis and mesophyll respectively^27,26^. While the *phdef* mutant flowers display a homeotic conversion of petals into sepals, the layer-specific *phdef* mutation results in two contrasted phenotypes^26^ (Figure 1A): flowers with a *phdef* mutant epidermis (that we named *star*) form a normal tube but small, star-shaped and unpigmented limb. Meanwhile, flowers with a *phdef* mutant mesophyll (that we named *wico* for *<u>wi</u>de <u>co</u>rolla*) have a much shorter tube than normal, but the limb is normally-shaped and pigmented, although petals are often pink rather than red. These flowers derive from the excision of the transposable element *dTph1* (inserted in the first exon of the *PhDEF* gene and knocking-out its function) in a unique cell layer, as we previously described^26^. The *star* and *wico* phenotypes show that PhDEF plays distinct roles in the two layers of the petal and drives partly independently limb and tube development. This material allows for the identification of cell layer-specific regulatory processes driven by PhDEF without the need to resort to tissue dissection. Here, to further understand how PhDEF achieves these cell layer-specific roles, we identified the cell layer-enriched target genes of PhDEF by characterizing layer-specific transcriptional profiles obtained by single-cell RNA sequencing (scRNA-Seq) and layer-specific binding profiles obtained by chromatin immunoprecipitation sequencing (ChIP-Seq), in wild-type (WT), *star* and *wico* petals. We found that, although *PhDEF* is similarly expressed in the two layers of the petal, PhDEF binds to and regulates the expression of many more target genes in the epidermis than in the mesophyll. The target genes of PhDEF display a combination of layer-specific and non-specific binding sites in their regulatory regions, suggesting a complex pattern of transcriptional control among cell layers. We also found differential enrichment of motifs under epidermal and mesophyll-specific peaks, suggestive of different protein partners for PhDEF in the two petal cell layers. This study offers a novel cell layer-specific perspective on transcriptional regulation and shows that homeotic regulators such as PhDEF are influenced by the pre-established cell layer identity.

**Figure 1.**
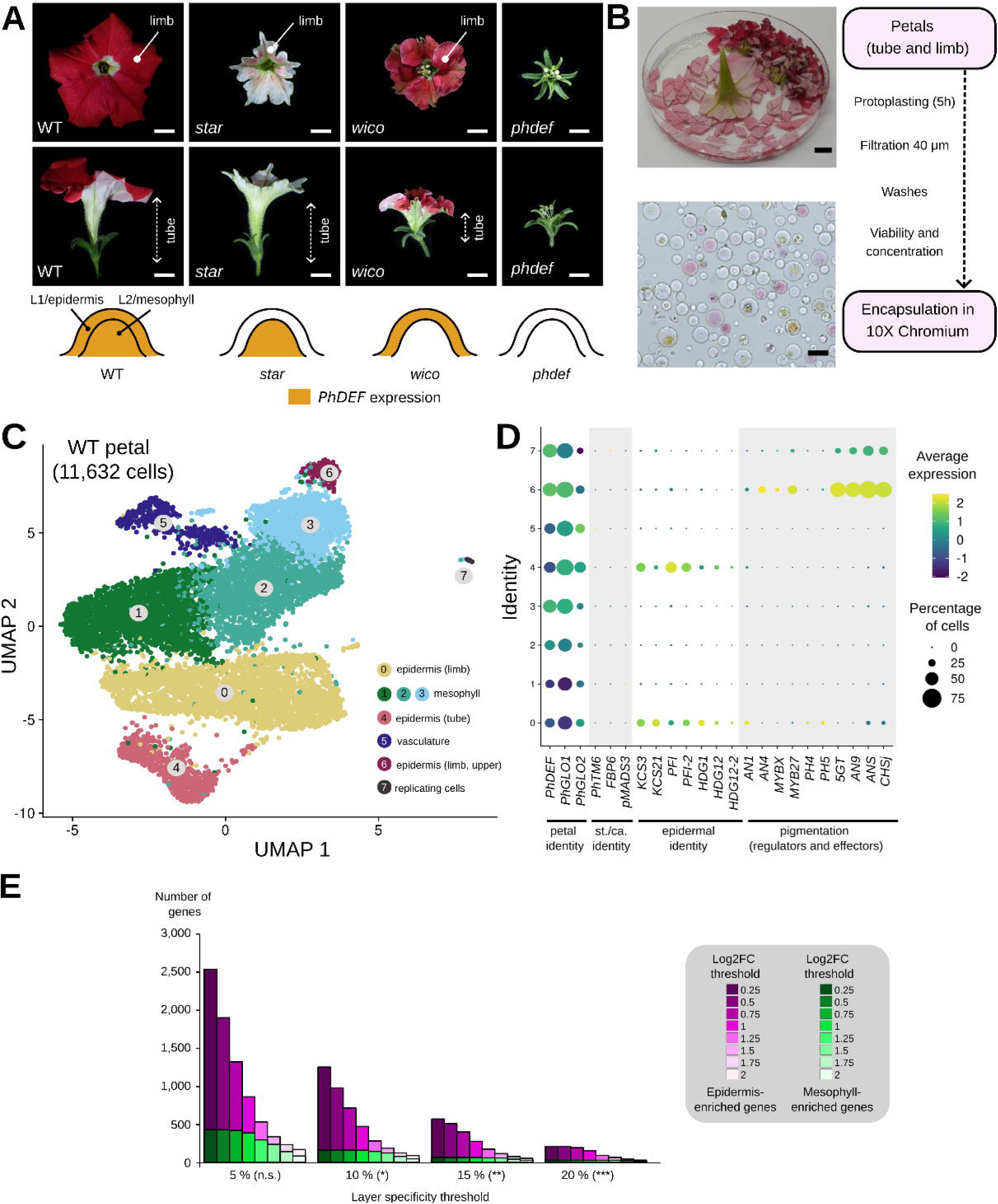
Single-cell RNA-sequencing separates epidermal and mesophyll cell types in the petunia petal. **(A)** Representative petunia wild-type (WT), *star*, *wico* and *phdef* flowers from an upper (top picture) and side view (bottom picture). The limb and tube are indicated. Scale bar = 1 cm. Below the flower pictures, a schematic petal primordium is depicted with L1 (future epidermis) and L2 (future mesophyll) cell layers, and *PhDEF* expression in the WT, *star*, *wico* and *phdef* primordia is indicated in orange. **(B)** Overview of the experimental protocol to produce petal protoplasts ready for isolation with the 10X Genomics Chromium device. Upper picture: a WT flower cut into small fragments in the cell wall-digesting solution (scale bar = 1 cm). Bottom picture: isolated protoplasts from a WT petal after a 5h-long digestion process, viewed by bright-light microscopy (scale bar = 50 µm). **(C)** Uniform Manifold Approximation and Projection (UMAP) plot of 11,632 WT petal cells sequenced for their transcriptome, after integration from two biological replicates. Clusters of cells are color-coded and are ordered from the biggest to the smallest, based on the number of cells. **(D)** Dotplot of key marker genes for petal identity, stamen and carpel identity, epidermal identity and pigmentation. The size of the dot represents the percentage of cells that express a given gene, and the color scale indicates the average expression level of the gene across all cells in a given cluster. st. = stamen, ca. = carpel. **(E)** Barplot of the number of genes enriched in the epidermis (magenta) or in the mesophyll (green), as defined by varying the log2FoldChange (log2FC) threshold and the layer specificity threshold (% of cells expressing the gene in the epidermis - % of cells expressing the gene in the mesophyll). The Kolmogorov-Smirnov two-sided test (KS test) was applied to compare the gene number distributions between epidermis- and mesophyll-enriched genes at a given layer-specificity threshold, n.s. non significant, * *p* < 0.05, ** *p* < 0.005, *** *p* < 0.001.

## RESULTS

### Single-cell RNA sequencing reveals separate clusters of epidermal and mesophyll cell types in the petunia petal

The *star* and *wico* phenotypes (Figure 1A) suggest that PhDEF regulates a distinct set of target genes in the two cell layers, contributing to either tube or limb development. We first aimed to identify the genes that exhibited enriched or non-enriched expression in the two WT petal layers, and for this we sequenced the transcriptome of single cells from wild-type (WT) petunia petals. To this end, the petals were isolated by carefully removing the stamen filaments that are fused to the base of the tube, and we performed a 5h-long protoplast isolation of the whole corolla (tube and limb) by cell wall-digesting enzymes (Figure 1B). While we were originally aiming to isolate protoplasts from petals at an early developmental stage (stage 8 as defined in^26^) to capture the dynamics of gene regulation during petal development, our protocol only allowed us to isolate a sufficient amount of protoplasts for scRNA-Seq in fully differentiated petals. A visual inspection of protoplasts under the microscope showed that the proportion of epidermal (pigmented with pink anthocyanins) and mesophyll (non-pigmented and/or chloroplastic) cells recovered was not significantly different to the proportion of epidermal and mesophyll cells observed in petal cross- sections (Figure S1), indicating that protoplasting allowed to efficiently recover cells from the two petal layers. Protoplasts were subjected to transcriptome sequencing with the 10X Genomics Chromium technology (Figure S2), resulting in the transcriptome from 11,632 cells from WT mature petals in two biological replicates (Figure 1C, Pearson’s correlation coefficient r = 0.96 between pseudo-bulk datasets of the two replicates).

The B-class petal identity genes *PhDEF*, *PhGLO1* and *PhGLO2* were found to be broadly expressed in all eight clusters (Figure 1D), while the stamen and carpel identity genes *PhTM6*, *FBP6* and *pMADS3*^12,28^ were hardly detected, indicating absence of contamination by stamen tissue. In order to assign identity to each cluster, we looked at known marker genes from the literature, identified their putative homologs in petunia (Table S1) and observed their expression in each cluster (Figure 1D, Figure S3). We also explored Gene Ontology (GO) term enrichment for the best marker genes of each cluster (Figure S4, Table S2) and we determined the limb or tube origin of respective clusters by RT-qPCR on a chosen set of cluster markers (Figure S5). The specific expression of the sucrose transporter SWEET11, and several UmamiT amino acid transporters and sulfate transporters, clearly marked cluster 5 as a group of vascular cells^29^ (Figure S3C). Similarly, the strong expression of several anthocyanin biosynthesis genes allowed the identification of cluster 6 as pigmented cells, hence cells from the upper limb epidermis^30^ (Figure 1D). Other known epidermal markers, such as ketoacyl-CoA synthases (KCS, wax synthesis enzymes)^31^ or members of the Homeodomain Glabrous (HDG) family^32^, identified clusters 0 and 4 as two other epidermal clusters (Figure 1D). Cluster 0 likely groups cells from the lower and upper limb epidermis; indeed, these cells faintly express pigmentation genes (Figure 1D); marker genes from this cluster are strongly expressed in limb tissue (Figure S5A); and *in situ* hybridization for the cluster marker *IAA14* indicated a clear expression in both upper and lower limb epidermis (Figure S5C). Consequently, this suggests that cells in cluster 6 represent only a fraction of the upper limb epidermal cells, *i.e.* the ones most strongly expressing pigmentation genes, perhaps the most distal cells from the limb that are the latest to differentiate. The highly specific expression of *TERPENE SYNTHASE1 (PhTPS1)*, a tube-specific gene involved in volatile emission inside the flower^33^, suggested that cluster 4 contains cells from the tube epidermis (Figure S5A, B). The strong and specific expression of several histone genes and S-phase genes (Figure S3B, D) identified the smallest cluster (7) as a cell-cycle state cluster (*i.e.* dividing cells, independent of a specific cell identity), as often found in single cell transcriptomics^34,35^. Finally, we assigned a mesophyll identity to the remaining clusters (1, 2 and 3), since they expressed genes strongly associated with photosynthetic activity for cluster 3 (Figure S3A) and with water transport for cluster 2 (Figure S4, Table S2), they did not express any of the epidermal or pigmentation markers (Figure 1D), and these clusters contained cells from either tube or limb origin (Figure S5A). We verified that the petunia homologs to genes defined as epidermal or mesophyll-enriched in tobacco petals also displayed a similar enrichment in our clusters^34^ (Figure S3E). Taken together, although there remains some uncertainty about the precise spatial location of each cluster, we are confident about their belonging to either the epidermis or the mesophyll layer.

Protoplasting is known to induce strong transcriptomic changes that could impact the clustering and the assignment of cell identities^37^. To estimate the effect of protoplasting on gene expression in our samples, we performed bulk RNA-Seq on protoplasted vs. intact WT petal tissue, and we observed a high correlation between the gene expression levels in the two conditions (r = 0.84). Still, a high number of differentially expressed genes (DEGs) were identified (5,571 and 5,891 genes respectively upregulated and downregulated by the protoplasting process, Table S3). We explored the effect of removing these DEGs on the clustering from the scRNA-Seq data (Figure S6): the UMAP plots exhibited a high degree of similarity in their shape irrespective of the removal of the DEGs, and the same clusters were recovered with only slight variation in the number of cells in each cluster. Therefore, we decided to keep all genes and all cells in our analysis.

In summary, our assignment of cluster identities revealed that the three mesophyll clusters are not clearly distinct in the UMAP space, in contrast to the three epidermal clusters that have distinct transcriptomic profiles. Importantly, there is a clear separation of epidermal and mesophyll cell types in the UMAP space, as previously observed in scRNA-Seq data from the petals of mature tobacco flowers^36^, showing that these layers have distinct transcriptomic signatures overall. Since our main interest was in the layer-specific functions of PhDEF, we next merged the three epidermal and the three mesophyll clusters together, setting apart the vasculature.

### The petal epidermis expresses more specific genes than the petal mesophyll

We first explored the difference in gene expression between WT petal layers. We computed gene differential expression between the epidermis and the mesophyll, and we examined the layer-enriched genes, *i.e.* with a higher expression level in one layer and with a high specificity of expression in one layer (higher percentage of cells expressing the gene in this layer). We noticed that there were consistently more genes whose expression is specifically enriched in the epidermis than in the mesophyll (Figure 1E). For instance, with log2FoldChange > 0.75 and layer-specificity of expression > 10 % (see Methods), there were 695 genes enriched in the epidermis compared to 151 genes enriched in the mesophyll (Table S4), and this asymmetry was observed at varying log2FoldChange and layer-specificity thresholds (Figure 1E). Some of the GO terms enriched in layer-enriched genes were consistent with the function of these layers (Figure S7, Table S2): the epidermis-enriched genes were associated with GO terms related to the metabolism of surface lipids or the phenylpropanoid pathway (an extensive network that leads to the production of lignin, anthocyanins and flavonols), while the mesophyll-enriched genes were associated with several GO terms related to photosynthesis. Overall, this initial analysis of layer-enriched genes showed that there are more epidermis-enriched genes than mesophyll-enriched genes in the WT petal.

### PhDEF regulates a different set of genes in the two petal layers

To explore how petal identity is defined by PhDEF in a layer-specific fashion, we generated scRNA-Seq data from 3,875 and 3,737 cells from *star* and *wico* mature petals respectively. Due to the difficulty in collecting high numbers of flowers from these genetic chimeras, we were only able to produce a single biological replicate for each. We integrated the WT, *star* and *wico* scRNA-Seq data using Harmony^38^, allowing for the identification of epidermal and mesophyll clusters in all samples (Figure 2A, B). The pseudo-bulk transcriptomes displayed a higher correlation between WT replicates (Pearson’s correlation coefficient, r = 0.96) than with the other genotypes (r = 0.9 between WT and *wico*, r = 0.93 between WT and *star*), as expected. The levels of expression of the main petal identity genes *PhDEF* and *PhGLO1* were consistent with our assumptions (Figure 2C-E): in WT petals, both genes were expressed evenly in all cell types, while in *star* petals we observed high expression in the mesophyll clusters and low expression in the epidermal clusters, and the converse was observed in *wico* petals. This is particularly apparent when looking at *PhGLO1* expression, that is higher than *PhDEF* expression at this stage (Figure 2D).

**Figure 2.**
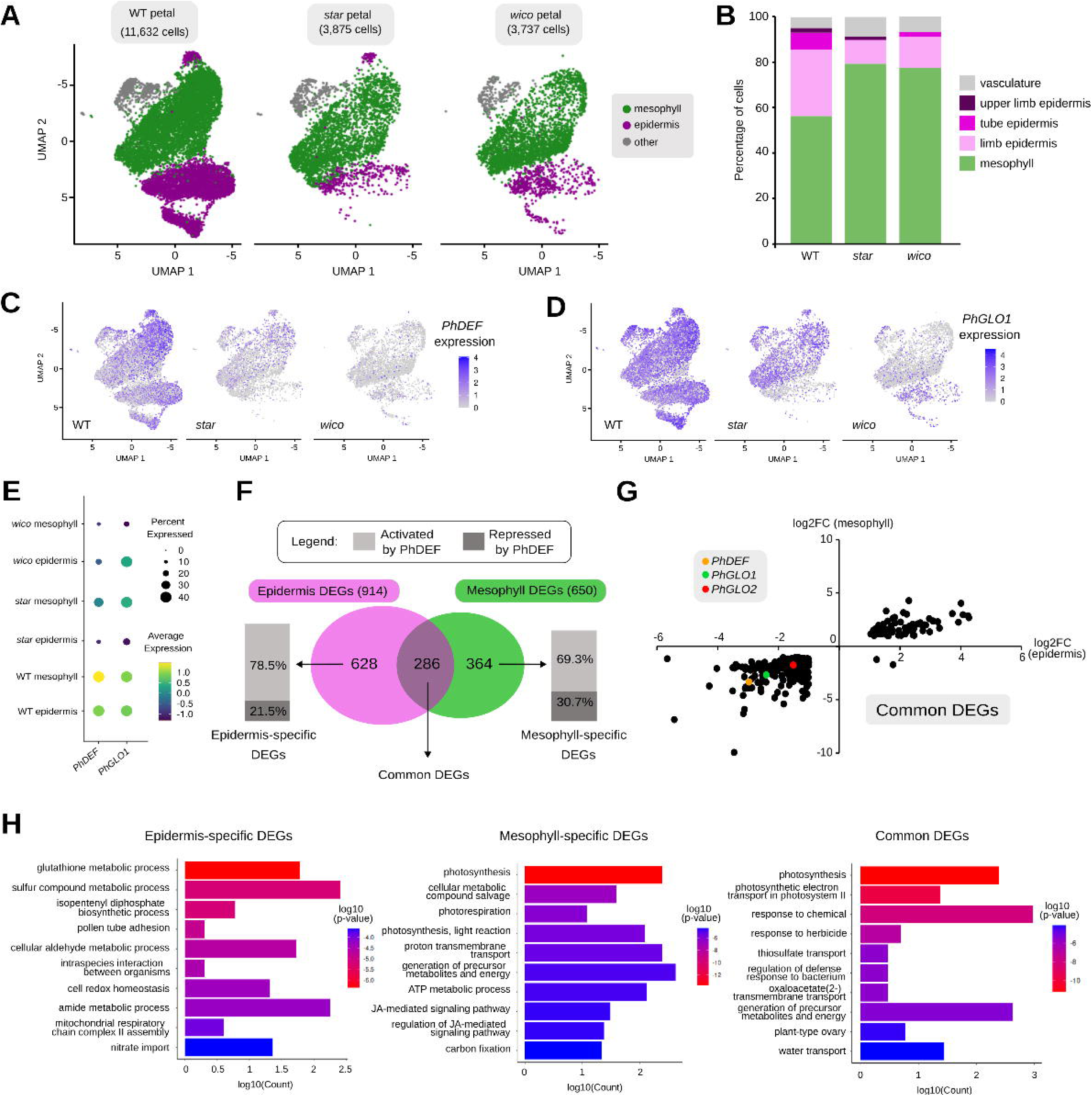
PhDEF regulates a different set of genes in the petal epidermis and mesophyll. **(A)** Uniform Manifold Approximation and Projection (UMAP) plots of 11,632 WT, 3,875 *star* and 3,737 *wico* petal cells sequenced for their transcriptome, after integration and clustering. Here, epidermal clusters (purple) and mesophyll clusters (green, without the vasculature) were merged. **(B)** Percentage of cells from the different epidermal and mesophyll identities identified from WT, *star* and *wico* petal scRNA-Seq. **(C)** Per-cell expression levels of *PhDEF* and *PhGLO1* in WT, *star* and *wico* displayed on UMAP plots. **(D)** Dotplot of PhDEF and PhGLO1 expression level (color- coded) and percentage of cells expressing the gene (coded in the size of the dot) in the epidermis and mesophyll cells of WT, *star* and *wico* petals. **(E)** Pearson’s correlation coefficient (r) plot between the pseudo-bulk transcriptomes from the mesophyll, epidermis and vasculature cells extracted from the WT, *star* and *wico* scRNA-Seq data. A Fisher r-to-z transformation test indicates that all correlation coefficients are significantly different (p < 0.05). **(F)** Venn diagram of the number of epidermis (purple) and mesophyll (green) DEGs, revealing the number of epidermis- and mesophyll-specific DEGs, as well as common DEGs (*i.e.* differentially expressed in both layers), based on WT, *star* and *wico* scRNA-Seq data. The percentage of activated (light grey) and repressed (dark grey) genes is also displayed. **(G)** Differential expression (log2FC = log2(FoldChange)) of the common DEGs in the epidermis or in the mesophyll, showing that almost all DEGs are either activated or repressed in both layers. Three genes of interest (*PhDEF*, *PhGLO1* and *PhGLO2*) are displayed as color points. **(H)** Ten most enriched Gene Ontology (GO) terms for biological processes in epidermis-specific (left), mesophyll-specific (middle) and common (right) DEGs, after GO term redundancy reduction and sorting by *p*-value.

When comparing the *star* and *wico* petals with WT petals, we did not uncover new clusters, which one might have expected from altered cell identity, but we observed changes in the proportions of cells in each cluster (Figure 2B). First, in *star*, a marginal number of upper limb epidermal cells were recovered, which corresponds to the small secondary L1-revertant sectors that are frequently observed in *star* petals^26^ (Figure S8) and that were thus later removed from the dataset. Apart from that, two major epidermal clusters (tube epidermis and limb epidermis) were strikingly depleted in cell numbers in *star* as compared to their WT counterparts. We interpret this as the result of two processes: (1) the lower proportion of limb tissue, that is mostly constituted of epidermal cells^39,26^, in *star* petals than in WT petals; (2) the alteration of epidermal cell identity by the *phdef* mutation, which might cause epidermal cells to cluster with mesophyll cells. In line with the latter, the pseudo-bulk transcriptome of the *star* epidermis was slightly more similar to the one from the WT mesophyll (r = 0.92) than to the one from the WT epidermis (r = 0.89) (Figure S9).

In *wico* petals, the main epidermal clusters were recovered in proportions higher than in *star* petals, but still much lower than in WT petals (Figure 2B). Surprisingly, the small cluster of strongly pigmented limb epidermal cells, which has a very clear transcriptional signature in WT petals (cluster 6 in Figure 1C), was not found in *wico* flowers. We speculate that this is because the *wico* flowers have a reduced pigmentation as compared to WT petals (Figure 1A, Figure S8), which we previously found to be caused by the 6-bp insertion left by the transposon excision from the *PhDEF* locus, slightly altering the function of PhDEF in activating pigmentation^26^. Consistently, we found 13 genes, out of the 42 associated with anthocyanin biosynthesis and its regulation, to be differentially expressed in the *wico* epidermis as compared to the WT epidermis. The depletion in epidermal cells in the *wico* UMAP space could therefore result from this slightly altered epidermal identity. In line with that, the pseudo-bulk transcriptome of the *wico* epidermis was as similar to the one from the WT epidermis (r = 0.9) than from the WT mesophyll (r = 0.9) (Figure S9). In contrast, in both *star* and *wico*, the mesophyll resembled more the WT mesophyll (r = 0.94 and 0.92 respectively) than the WT epidermis (r = 0.88 and 0.84 respectively) (Figure S9). This global inspection of layer-specific *star* and *wico* transcriptomes suggested that alteration of epidermal identity tends to shift the transcriptome towards a mesophyll-like transcriptome, while alteration of mesophyll identity did not have the converse effect; therefore, the mesophyll identity appears as the default cell identity in the petal.

Next, we explored in detail the effect of *phdef* mutation on gene expression in a given petal layer. Since the *star* flowers are only mutant for *PhDEF* in the epidermis, we defined epidermal- specific DEGs as differentially expressed in the *star* epidermis as compared to the WT epidermis (with |log2FC| > 1, adjusted *p-*value < 0.01 and specificity of expression >10 %, see Methods). A major limitation of this definition is that, as previously shown, *star* epidermal cells might cluster with mesophyll cells because their transcriptional identity has been altered. This would lead to an artificial depletion of *star* epidermal cells in the analysis, which would result in the underestimation of epidermal-specific DEGs. Conversely, mesophyll-specific DEGs were defined as the genes differentially expressed in the *wico* mesophyll as compared to the WT mesophyll. We found that 286 genes were DEGs in both layers, while 628 were exclusive DEGs in the epidermis and 364 were exclusive DEGs in the mesophyll (Figure 2F, Table S4). Hence, even accounting for a possible underestimation of epidermal-specific DEGs, we found more epidermis-specific DEGs than mesophyll-specific DEGs. These genes were either activated (75%) or repressed (25%) by PhDEF (Figure 2F), and there was a slightly higher proportion of genes whose expression was activated by PhDEF in the epidermis (78.5%) than in the mesophyll (69.3%, Figure 2F). The vast majority (99.3%) of the genes commonly regulated by PhDEF in both layers, hereafter named common DEGs, were regulated in the same direction in the two layers, *i.e.* either activated (75.2% of genes) or repressed (24.8% of genes, Figure 2G). Hence, the direction of regulation by PhDEF is not heavily influenced by the petal layer in which it acts. As expected, *PhDEF*, *PhGLO1* and *PhGLO2* were found among the 287 common DEGs, activated by PhDEF in the two petal layers (Figure 2G, Table S4). Gene Ontology (GO) terms enriched in epidermis-specific DEGs were related to redox processes (glutathione metabolism, cell redox homeostasis), while those enriched in mesophyll-specific DEGs were strongly associated with photosynthesis, and various different GO terms were enriched in common DEGs (Figure 2H, Table S2).

In summary, our scRNA-Seq dataset from layer-specific *phdef* mutant flowers indicates that PhDEF regulatory action is largely different between the two cell layers, as the number of common DEGs is lower than the number of layer-specific DEGs, and this is consistent with the drastically different *star* and *wico* phenotypes. We also found about two times more epidermal-specific than mesophyll-specific DEGs.

### PhDEF binds to more target genes in the petal epidermis than in the petal mesophyll

The scRNA-Seq transcriptomic data confounds the effect of direct gene regulation by PhDEF, and the indirect gene regulation that results from it, which might be extremely different in terms of numbers of genes, direction and intensity of the transcriptional regulation among the pathways affected. To circumvent this limitation, we then attempted to identify PhDEF direct targets in the two petal layers. For this, we performed chromatin immunoprecipitation followed by sequencing (ChIP-Seq) in WT, *star* and *wico* petals, using a custom antibody directed against the endogenous PhDEF protein^26^. After a set of preliminary tests to evaluate ChIP efficiency (Figure S10 and Methods), we performed the experiment on two biological replicates, on flowers at stage 8 when the tube is about half its final size, in an attempt to capture most gene regulation events underlying both limb and tube development^26^. After sequencing and mapping the reads on the *P. axillaris* genome^30^, we called the peaks for each replicate separately, then computed the Irreproducible Discovery Rate (IDR)^40^ to identify reproducible peaks and removed peaks detected in the input chromatin (see Methods), which resulted in 3,134, 4,276 and 2,216 peaks for WT, *wico* and *star* respectively (Table S5, Table S6). We observed that PhDEF bound to similar regions of the genome in WT, *wico* and *star*, with a generally higher read coverage in WT and *wico* samples than in *star* samples (Figure 3A, B, Table S6). Importantly, this is consistent across the two ChIP replicates for each genotype (Table S5), and is not due to a more variable PhDEF expression in the mesophyll as compared to the epidermis (Figure S10), that could have caused a general lower enrichment of PhDEF binding in the mesophyll. This indicates that binding of PhDEF to the genome is more frequent in the epidermis than in the mesophyll, but overall takes place in the same regions of the genome.

**Figure 3.**
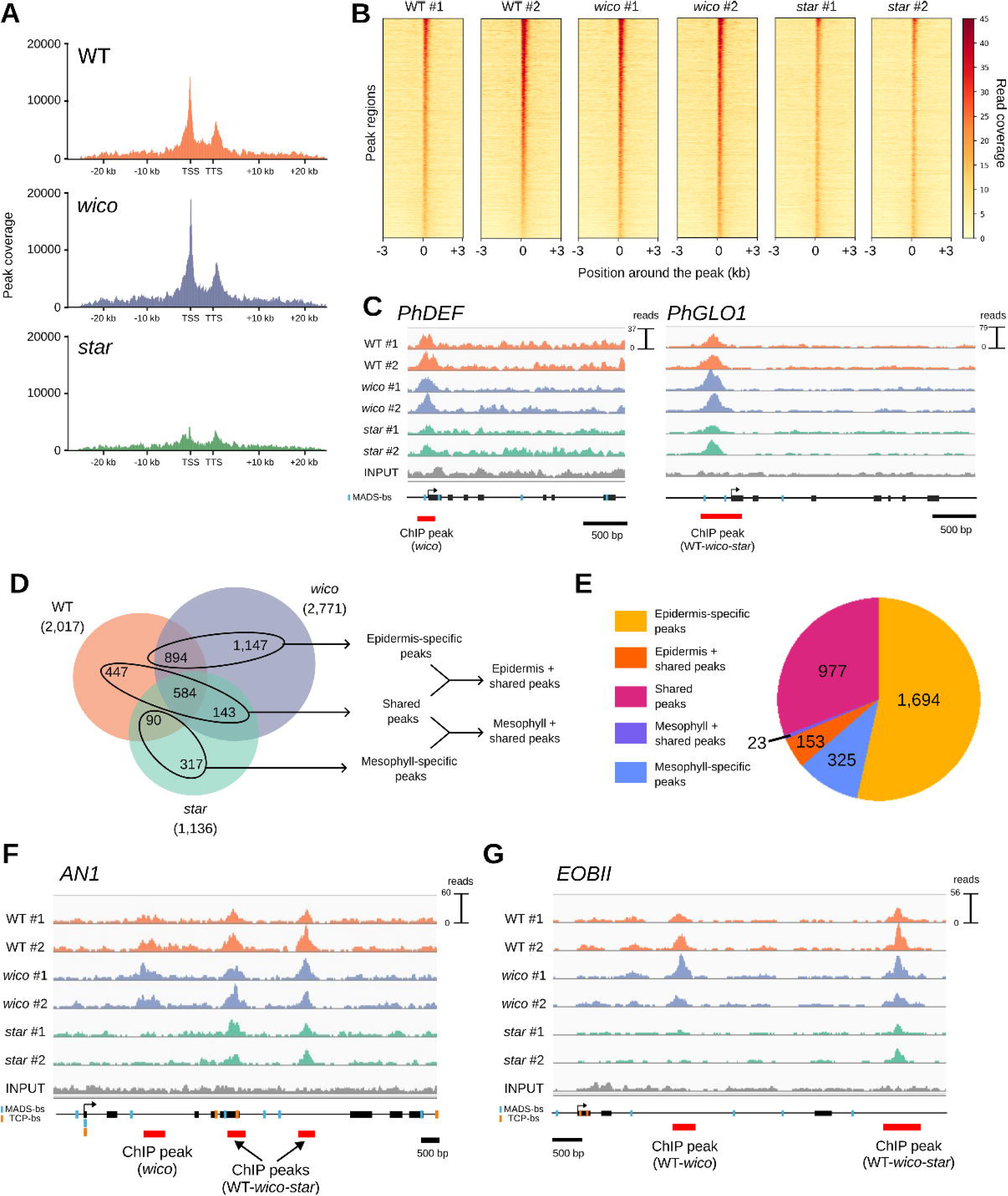
PhDEF binds to more target genes in the epidermis than in the mesophyll. **(A)** Metaplot of PhDEF binding sites in WT, *wico* and *star* petals, with the distance from the Transcription Start Site (TSS) and the Transcription Termination Site (TTS). **(B)** Heatmap of the read coverage of all peaks detected in WT samples, and of corresponding regions of the genome in *wico* and *star* samples. The peaks are sorted according to the highest to lowest read coverage in WT #2, so that the same regions of the genome are in the same line in all librairies. The read coverage is color-coded, and position 0 represents the start of the peak. **(C)** Read coverage for PhDEF binding to the genomic regions of *PhDEF* and *PhGLO1* in WT, *star* and *wico* petals. Peaks that pass our pipeline for reproducibility (see Methods) are displayed as thick red lines. Predicted MADS-binding sites (MADS-bs) are indicated by blue lines, while no TCP-bs were found in these sequences. Although the peak in the promoter region of *PhDEF* does not pass the IDR threshold, its read coverage profile strongly suggests that PhDEF binds to this position. **(D)** Intersection of the number of gene-associated peaks from WT, *star* and *wico* ChIP-Seq data, and definition of the possible PhDEF binding profiles. The intersection occasionally resulted in the artifical duplication of peaks, and once corrected, this marginally changed the total number of peaks identified for each genotype in the intersection. **(E)** Distribution of PhDEF binding profiles across its target genes. **(F-G)** PhDEF binding profile over the genomic locus of AN1 (**F**) and EOBII (**G**). Predicted MADS-binding sites (MADS-bs) are indicated by blue lines, and predicted TCP binding sites (TCP-bs) are indicated by orange lines.

In total 2,017, 2,771 and 1,136 peaks were associated with genes for WT, *wico* and *star* respectively (Table S6, Table S7). Most of these peaks were located in promoter regions (defined as 6 kb upstream of the start codon, 34.5% for WT, as compared to a random 14.9% occupancy, *p* = 2.06e^-153^, hypergeometric test) or terminator regions (defined as 6 kb downstream of the stop codon, 20.1% for WT, *p* = 6.44e^-13^) of genes, with a small and non-enriched fraction found in gene bodies (9.8% on average, *p* = 0.51) (Figure 3A, Table S6). These proportions were similar in WT and *wico*, but *star* displayed a lower proportion of peaks located in the promoter regions (24.5% for *star* vs. 34.5% and 36.2% for WT and *wico*, respectively, Table S6), suggesting that binding of PhDEF to promoter regions is more frequent in the epidermis than in the mesophyll. Out of the genes bound by PhDEF in WT, we found a substantial proportion (37.0%, *p* = 1.20e^-118^, hypergeometric test) also differentially expressed in the previously published *phdef-151* mutant transcriptome^26^, which is consistent with the proportions that have been reported from previous ChIP-Seq assays on MADS-box proteins^23,41–43^. In particular, we could detect clear binding of PhDEF on its own promoter, and on the promoter of the other B-class gene *PhGLO1* (Figure 3C), that are two established regulatory targets of PhDEF, detected as differentially expressed both in bulk RNA-Seq and in scRNA-Seq.

We next examined the intersection of gene-associated peaks (including promoter, gene body and terminator regions) between WT, *wico* and *star* flowers (Figure 3D, Table S7). Strikingly, and consistently with the high number of peaks detected in *wico*, there were many more epidermal- enriched peaks (*i.e.* found in WT+*wico* or *wico* alone, 2,041 peaks in 1,896 genes) than mesophyll-enriched peaks (*i.e.* found in WT+*star* or *star* alone, 407 peaks in 398 genes). A large number of peaks (1,176 in 1,110 genes) were also shared between layers (*i.e.* found in WT, *star+wico* or *WT+star+wico*). When shared, most peaks were higher in *wico* than in WT than in *star* (see Figure 3C for *PhGLO1* for instance), supporting again that the binding of PhDEF to its target sites is more frequent in the epidermis than in the mesophyll. Hence for most peaks, PhDEF binding in the WT sample can be interpreted as the average of frequent PhDEF binding in the epidermis and rare or absent PhDEF binding in the mesophyll, explaining why some peaks are detected in *wico* but not in WT. The 317 peaks uniquely found in *star* were also weakly bound in the epidermis, and the 1,147 peaks uniquely found in *wico* were also weakly bound in the mesophyll (Figure S11), showing that differential binding between layers is quantitative, with a relative enrichment in the epidermis or in the mesophyll that can differ across genomic regions.

While most of PhDEF target genes (91.8% on average) were associated with a single ChIP peak, others displayed between 2 and 4 binding sites for PhDEF (Table S6). To our surprise, these different binding sites over a single target gene were not necessarily occupied by PhDEF similarly in the two petal layers: for instance, the gene encoding the key regulator of anthocyanin biosynthesis ANTHOCYANIN1 (AN1)^44,45^ displayed three binding sites for PhDEF in its regulatory regions, two of them being epidermal-specific and one of them being shared between petal layers (Figure 3F). A similar binding profile, with one epidermal-specific and one shared peak, was observed for the gene encoding the MYB-R2R3 transcription factor EMISSION OF BENZENOIDSII (EOBII), a major regulator of scent emission and petal maturation^46–48^ (Figure 3G). These complex binding profiles accounted for a rather small number of genes (153 genes with epidermis-specific + shared peaks, 23 genes with mesophyll-specific + shared peaks), compared to the vast number of genes displaying a single type of PhDEF binding profile (1,694 genes with only epidermis-specific peaks, 977 genes with only shared peaks and 325 genes with only mesophyll-specific peaks). Hence, the PhDEF binding profile can be complex, with layer-specific or -aspecific binding sites, which might reflect differential transcriptional regulation between cell layers.

### Epidermis DEGs display more epidermal and shared binding sites for PhDEF

We next wanted to explore further how different PhDEF binding profiles are associated with the expression profiles of its target genes by combining our transcriptomic and ChIP-Seq assays on WT, *star* and *wico* flowers. We compared our ChIP-Seq results with either the scRNA-Seq dataset, or with the bulk RNA-Seq dataset on WT, *star* and *wico* petals that we previously obtained^26^. These two datasets have complementary benefits: the scRNA-Seq dataset provides a layer-specific view of gene differential regulation, while the bulk RNA-Seq dataset has been generated at the exact same stage as ChIP-Seq and better captures lowly expressed genes. Both bulk RNA-Seq and scRNA-Seq datasets overlap with the ChIP-Seq dataset more than by chance (Figure S12, *p* = 4.73e^-13^ and *p* = 1.37e^-04^ respectively, hypergeometric test).

Using bulk RNA-Seq data, we explored the distribution of PhDEF binding profiles for DEGs in *wico* and/or in *star* petals at stage 8 (Figure 4A), and compared it with the distribution we had previously computed for all PhDEF target genes (Figure 3E). Overall, distributions were very similar, with a majority of epidermis-specific peaks and shared peaks, as previously observed. A significantly different distribution of PhDEF binding profiles was observed for *star*-specific DEGs (Figure 4A, Chi^2^ goodness-of-fit, *p* = 0.039), with more genes displaying epidermis+shared binding sites (test of equal proportions, one-sided, *p* = 0.037). No significant deviation from the expected distribution is found for *wico*-specific DEGs (Chi^2^ goodness-of-fit, *p* = 0.080) nor for common DEGs (*p* = 0.064), but *wico*-specific DEGs were enriched for mesophyll-specific peaks (test of equal proportions, one-sided, *p* = 0.024). To explore whether layer-specific regulation profiles might be blured in the bulk RNA-Seq data, since it mixes the transcriptome of both cell layers, we analyzed PhDEF binding profile in the layer-specific DEGs identified by scRNA-Seq (Figure 4B). Again, distributions of PhDEF binding profiles were highly similar for epidermal-specific, mesophyll-specific or shared DEGs, (Figure 4B, Chi^2^ goodness-of-fit, *p* = 0.079, 0.28 and 0.94 respectively), but we could find an enrichment of epidermis+shared binding sites in epidermis- specific DEGs (test of equal proportions, one-sided, *p* = 0.033). Overall, and keeping in mind the limitations of these analyses, these results show that epidermis DEGs tend to display more epidermis-specific and shared binding sites for PhDEF, suggesting that layer-specific binding and transcriptional regulation are partially linked, at least in the epidermis.

**Figure 4.**
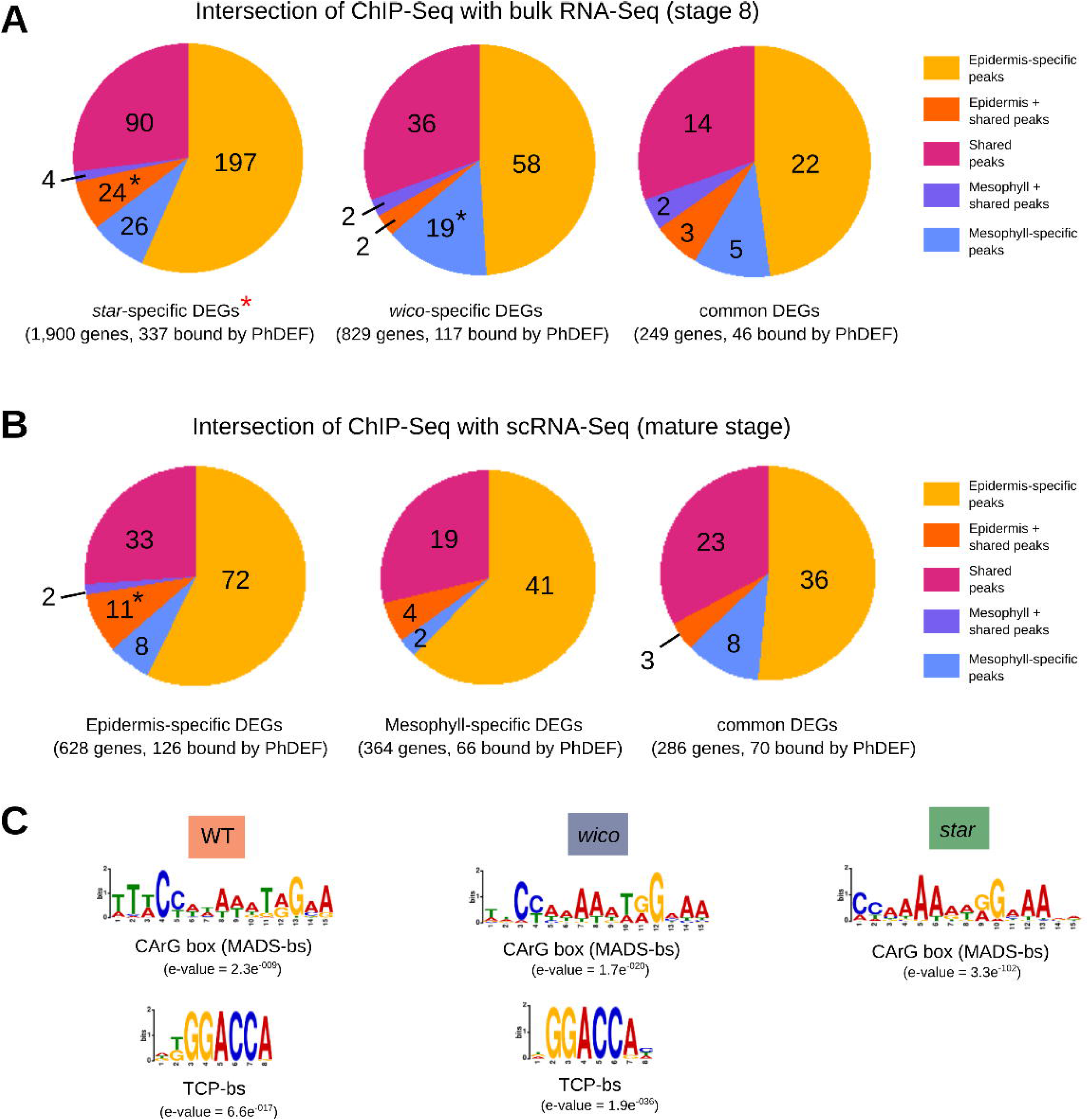
Intersection between PhDEF binding and regulatory profiles, and motif enrichment under PhDEF binding sites. **(A)** Distribution of PhDEF binding profiles across genes differentially expressed (DEGs) in *star* only, *wico* only, or both in *star* and *wico* (common DEGs), as compared to WT, from bulk RNA-Seq of petals at stage 8. Stars indicate a significant deviation from expected PhDEF binding profiles, either for the whole distribution (red star, Chi2 goodness-of-fit test) or an enrichment of individual categories (black star, test of equal proportions, one-sided). Binding categories with less than 5 genes were not tested for significant enrichment. **(B)** Distribution of PhDEF binding profiles across epidermis-specific, mesophyll-specific or common (in both layers) DEGs, obtained from WT, *star* and *wico* scRNA-Seq by comparing layer-specific transcriptomes. **(C)** Selected motifs enriched under PhDEF ChIP-Seq peaks for WT, *wico* and *star* petals, bs = binding site. The full list of motifs detected is in Figure S9.

### Different motifs are enriched under layer-specific PhDEF binding sites

A possible mechanism by which PhDEF regulates different target genes in the two petal cell layers could be its interaction with different protein partners, which would modify its DNA-binding specificity. This should be readily detectable in the enriched motifs for transcription factor binding under ChIP-Seq peaks. When examining these motifs, we found that gene-associated ChIP-Seq peaks for WT, *wico* and *star* were all enriched in CArG-boxes (Figure 4C, Figure S13), the typical MADS-box binding site. This likely reflects PhDEF direct binding to these regions but might also be the result of other MADS-box proteins binding, together with PhDEF as dimers or tetramers. Indeed, PhDEF is known to interact with the B-class proteins PhGLO1 and PhGLO2^11^, and with the E-class proteins FBP2 and FBP5 (orthologous to Arabidopsis SEPALLATA3 (SEP3) and SEP1/2, respectively)^21,22^. We examined the cell layer-specific expression of these genes in our WT scRNA- Seq dataset and found that the B- and E-class genes were overall homogeneously expressed between the two petal layers or hardly detected (Figure S13A). It is therefore unlikely that different floral quartets would form in the epidermis and in the mesophyll. Apart from CArG boxes, we also found enrichment for TCP (TEOSINTE BRANCHED 1 / CYCLOIDEA / PROLIFERATING CELL NUCLEAR ANTIGEN FACTOR1) binding sites in WT and *wico* gene-associated peaks, but not in *star* (Figure 4C, Figure S12). TCP transcription factors are known to be partners of MADS-box proteins^49,50^, and also to regulate the same genes as the ones regulated by MADS-box proteins^51^, explaining why their motifs are occasionally found in co-occurrence under ChIP peaks^43,52^. The fact that this motif is not found enriched under *star* peaks could be due to two reasons: either the number of peaks in *star* was too low to actually detect such enrichment; or TCP proteins could be partners of PhDEF or coregulators of PhDEF target genes only in the petal epidermis. As a support to the second hypothesis, we detected a strong enrichment of predicted TCP binding sites under epidermal-specific peaks and shared peaks, but only a weak one under mesophyll-specific peaks, as compared to random sequences (Table S8). For gene-associated peaks, 25.2 % of epidermal-specific peaks, and 25.0 % of shared peaks, had at least one predicted TCP binding site, in contrast to only 11.8 % of mesophyll-specific peaks (Table S8). We also examined the expression of petunia *TCP* genes^53^ in our WT petal scRNA-Seq dataset and found a clear enrichment for *PaTCP4a* in the epidermis, while other *TCP* genes displayed low expression and/or similar expression in the two cell layers (Figure S14B). The expression of this gene increases during petal development, is lower in *star* than in *wico* flowers at stage 8 (Figure S14C), and is also higher in the *star* and *wico* epidermis than in their corresponding mesophyll tissue (Figure S14D). The expression of *PaTCP4a* was also reported to be higher in *Petunia axillaris* flowers with large petals than in flowers with small petals^53^. PaTCP4a therefore appears as a good candidate to be an epidermal-specific partner of PhDEF in the petal, which remains to be verified experimentally.

## DISCUSSION

In this work, we investigated the cell layer-specific functions of the petal identity regulator PhDEF in petunia, based on the initial observation that epidermal- or mesophyll-specific expression of *PhDEF* yielded drastically different petal phenotypes. We found that PhDEF regulates a different set of genes in the two petal layers, with a major binding and regulatory action in the epidermis. We also identified target genes with complex binding profiles, and overall the association between binding and regulatory profile of PhDEF target genes was limited, but significant for genes bound and regulated in the epidermis, displaying more epidermal+shared peaks than expected. This suggests that PhDEF can achieve layer-specific regulation of its target genes through layer-specific binding, in particular in the epidermis. The reasons for the otherwise low association between binding and regulatory profiles are likely manifold: PhDEF binding does not always entail target gene regulation; PhDEF can act either as an activator or as a repressor of gene transcription; long- distance looping between PhDEF binding sites (as has been shown for MADS-box proteins^51,52^) creates complex binding profiles that exceed the environment of a single gene; and most importantly, many other transcription factors than PhDEF regulate the expression of its target genes.

Globally, gene transcriptional regulation was consistently more prominent in the epidermis than in the mesophyll. First, we observed that the epidermis expresses more specific genes than the mesophyll, which, to our knowledge, had not been reported before. Hence, the mesophyll might be regarded as a default identity for the petal cells, upon which the regulation of a large set of genes is appended for the specification of epidermal identity. Second, we found that *phdef* mutation leads to the differential expression of more genes in the epidermis than in the mesophyll. Third, we observed that PhDEF strikingly binds to more binding sites and target genes in the epidermis than in the mesophyll. This prominent role of PhDEF in the petal epidermis aligns with the multiple specific features found in this tissue, and in epidermal tissues in general: trichomes, stomatas, glandular cells or conical cells; and the very specific functions that the epidermis can fulfil for pigmentation, volatile emission, cuticle biosynthesis or defense against biotic and abiotic stress^56–59^. In contrast, the mesophyll appears less differentiated, with cells visually quite similar and fulfilling less diverse functions than the epidermis^57,60,61^, although this might be biased by the low accessibility of this tissue, and the reduced interest it has attracted. Several epidermal-specific genes have been described over the years, in particular the early specifiers of embryonic epidermal identity *MERISTEM LAYER1* (*ML1*) or *PROTODERMAL FACTOR2* (*PDF2*) from the previously cited family of HDG transcription factors^62–65^, or the genes acting in later epidermal-specific functions, such as stomata^59^ or trichome^66^ formation, cuticle synthesis^67^ or pigmentation^68^. In contrast, a much smaller number of genes specific to the L2 layer have been described^64,69^, and these genes are all expressed post-embryonically^64^, suggesting the possibility that the sub-epidermal identity does not require specific regulators for its determination. Therefore, establishing epidermal identity might truly require the regulation of several specific genes, starting from the canvas of a default mesophyll identity. Linking this prominent action of PhDEF in the epidermis with the *star* and *wico* phenotypes, suggests that building the limb, that is mostly constituted of epidermal tissue, requires the regulation of a larger set of genes than building the tube.

We have two main hypotheses explaining why PhDEF adopts a different binding and regulatory behaviour in the epidermis and in the mesophyll: First, PhDEF might have access to different protein partners in the two petal layers, and since binding to different protein partners entails changes in DNA-binding specificity, which has been demonstrated for MADS-box proteins^70,71^, this would lead to the regulation of different target genes. Our identification of a TCP binding motif enriched under epidermal-specific PhDEF binding sites suggests that MADS-TCP complexes could contribute to the specific regulation of PhDEF epidermal targets. However, it would account for 25% of PhDEF target sites at most. Second, PhDEF might have access to different target genes in the two petal layers, due to different chromatin accessibilities. Recently, single nuclei ATAC-Seq (Assay for Transposase-Accessible Chromatin) assays in maize and sorghum leaves revealed that epidermal and mesophyll chromatin accessibilities were indeed very different, with more accessible regions in the epidermis than in the mesophyll in both species^72^. MADS-box proteins have also been shown to interact with chromatin remodellers^73,74^, which might participate in the reinforcement of differential chromatin accessibility between layers. The fact that the difference between epidermal and mesophyll binding of PhDEF is mostly quantitative, with a more frequent binding in the epidermis, seems in favour of an elevated chromatin accessibility in the epidermis. Apart from these two main non-mutually exclusive scenarios, a major aspect that remains unexplored in our study is the presence and activity of the PhDEF protein in the petal cell layers. The mere enrichment of PhDEF in the epidermis could explain its prominent regulatory role in this layer, which would then open investigations on the regulation of *PhDEF* translation, PhDEF protein movement between layers^75,76^, or PhDEF protein degradation.

Overall, our study provides a cell-layer-specific view on transcriptional regulation, directly accessible here thanks to the *star* and *wico* periclinal chimeras. Other studies than ours have reported that regulators could have different functions in different cell layers^77–82^, for other floral homeotic genes but not exclusively, which suggests that cell layer identity likely impacts transcription factor activity as a general rule.

## ACKNOWLEDGMENTS

We thank Francesco Quattrocchio and Shuangjiang Li for sharing their protocol for protoplast isolation; Cyril Dégletagne and the Cancer Genomics Platform from the Centre de Recherche en Cancérologie de Lyon for support in the scRNA-Seq experiment; Nicolas Dalle and Annick Dubois for assistance in chromatin immunoprecipitation; Benjamin Gillet and Sandrine Hugues f rom the sequencing platform of the Institut de Génomique Fonctionnelle de Lyon for library preparation and sequencing of the transcriptomes of this study; the Master of Biology of the Ecole Normale Superieure de Lyon for their participation in funding and analyzing the ChIP-seq experiment; and Sergio Sarnataro from Spatial-Cell-ID for advice on scRNA-Seq analysis. We also thank three anonymous reviewers for their constructive criticism that greatly helped improve this manuscript. This work was supported by a grant to QC-S and MM from the Agence Nationale de la Recherche (grant ANR-19-CE13-0019, FLOWER LAYER), by a grant to ED from the French Ministry of Higher Education and Research, by grants to DB from the Agence Nationale de la Recherche (grants ANR-21-CE12-0036-01 and ANR-21-CE20-0007-02) and by the EquipEx+ Spatial-Cell-ID under the “Investissements d’avenir” program (ANR-21-ESRE-00016). We gratefully acknowledge support from the CBPsmn (PSMN, Pôle Scientifique de Modélisation Numérique) of the ENS de Lyon for the computing resources. The platform operates the SIDUS solution^83^ developed by Emmanuel Quemener.

## AUTHOR CONTRIBUTIONS

QC-S, ED, DB and MM conceived the project and designed experiments. QC-S, ED, PC, PM, and SRB performed the experiments and QC-S, ED, ED-B, BL, CR, JJ and MM analyzed the data. QC-S, ED, DB and MM designed figures and wrote the manuscript, with input from all authors.

## MATERIAL AND METHODS

### Plant material and culture conditions

Plants were grown in a culture room in long day conditions (16h day at 22°C, 8h night at 18°C, 75-Valoya NS12 LED bars, light intensity: 130 μE, 60% humidity). All plant material derives from the *Petunia x hybrida* R27 line containing the active *dTph1* transposable element. The *wico* and *star* flowers spontaneously arose from *phdef-151* homozygous mother plants, by the cell layer-specific excision of *dTph1*, resulting in branches carrying either *wico* or *star* flowers that were repeatedly obtained from several different *phdef-151* individuals, as described in^26^. These branches were subsequently maintained by cuttings.

### Histology

Petal cross-sections and toluidine blue staining were performed as described in^26^. Cell types were counted as indicated in Figure S1, from 4 different cross-section images per limb and per tube (n = 1,038 limb cells and 2,736 tube cells counted in total).

### Bulk RNA-Seq

From WT petals at stage 12^26,39^, a fraction of the corollas was directly flash-frozen; the rest was protoplasted as described below and the protoplast pellet was flash-frozen. RNA extraction was performed with Sigma’s Spectrum™ Plant Total RNA Kit with On-Column DNase I Digestion, following manufacturer’s recommendations. RNA integrity and quantity were determined using a Bioanalyzer RNA 6000 Nano assay (Agilent), libraries were prepared with the CORALL RNA-Seq Library Prep (Lexogen) and sequenced with an Illumina NextSeq500 (single-end reads, 84 bp). Reads were mapped on the *P. axillaris* transcriptome^84^ as described in^26^. Differential gene expression was computed with DESeq2^85^ (1.34.0) in RStudio (R 4.4.1, RStudio 2024.04.2).

### RT-qPCR

For RNA extraction, limb or tube tissue was collected from WT flowers at anthesis in 3 biological replicates. For limb samples, the limb from 1 petal from 3 different flowers constitutes one replicate; for tube tissue, the whole tube from 3 different flowers constitutes one replicate. RNA extraction, Reverse Transcription (RT), quantitative PCR (qPCR) and result analysis were performed as described previously^22^. Primer sequences are available in Table S9.

### *In situ* hybridization

*In situ* hybridization was performed as previously described^26^, using *f3h* mutant flowers with white petals^86^, to avoid detecting an aspecific signal in pigmented epidermal cells. The primers used to synthesize the antisense probe for *IAA14* can be found in Table S9.

### Single-cell RNA-Seq

#### Petal protoplast isolation

Protoplast isolation was performed in sterile conditions. WT, *star* and *wico* corollas (ca. 15, 30 and 30 flowers per replicate, respectively) were briefly soaked in 70% ethanol and in 0.5% bleach for 30’’ then rinsed 3 times with sterile water. Sterile corollas were transferred in a petri dish containing 2 mL of Digestion Mix (0.4% macerozyme R-10, 0.8% Cellulase Onozuka R-10, w/v in TEX Buffer (3.1 g/L Gamborg B5 salts, 500 mg/L MES, 750 mg/L CaCl_2_*2H_2_O, 250 mg/L NH_4_NO_3_, 136.9 g/L Sucrose, pH 5.7)) and cut in ca. 0.5 cmZ pieces, using a new scalpel blade for each corolla to reduce tissue wounding. 10 mL of Digestion Mix were added and digestion was performed for 5 h at 26°C in the dark, with gentle orbital agitation (20 rpm) for the last 15’. After filtration through a 40 µm mesh, the volume was adjusted to 25 mL using 0.4 M Sucrose (492 mOsm.kg-1 H_2_O) and the mixture was centrifuged for 10’ at 100 g with acceleration 2/9 and deceleration 0/9 with a swing-out rotor. The underlying buffer was removed using a peristaltic pump (Gilson MINIPULS™ Evolution) connected to a sterile Pasteur pipette without perturbing the protoplast layer, at a rate of ca. 100 µL/sec. After adjusting again the volume to 25 mL using 0.4 M Sucrose, the whole process was repeated twice. Protoplast concentration and viability was assessed using a Kova slide and staining with 1% Evans Blue dye solution (w/v in 0.4 M Sucrose) as described in^87^. Cell types were estimated as indicated in Figure S1, from 4 different pictures taken from a single protoplast isolation event (n = 3,338 cells counted in total).

#### From single cell transcriptome sequencing to clustering

Protoplast suspensions were adjusted to 345, 480, 590 and 560 cells/µL with Mannitol-BSA (0.44 M Mannitol, 0.1 % BSA (w/v), 498 mOsm.kg-1 H2O) for WT (first replicate), *star*, *wico* and WT (second replicate) respectively. Protoplasts were loaded in the 10x Genomics Chromium chip and libraries were prepared using the Chromium Next GEM Single Cell 3ʹ Reagent Kits v3.1 (dual index) kit following manufacturer’s instructions. Libraries were sequenced on an Illumina NovaSeq 6000 SP or S1. Read quality was checked with FastQC^88^ (v0.12.1) and reads were aligned on the *P. axillaris* v1.6.2 transcriptome (annotation from^30^ transferred to a genome improved by HiC^26,89^, which was the best available version of a Petunia genome at that time). CellRanger count (v7.0.1, 10X Genomics) was used to filter reads, count barcodes and UMIs, and generate the HDF5 matrix. The second WT replicate was split and sequenced in two technical replicates on the same Chromium chip, and read counts were aggregated with the Cell Ranger function aggr. Normalized gene expression, UMAP plots and cluster markers were computed with Seurat 5.1.0^90,91^ in R^92^ (v4.3.3) as follows, using default parameters unless stated otherwise: cells with less than 200 genes and genes dectected in less than 3 cells were removed; data was normalized with NormalizeData, highly variable features were identified with FindVariableFeatures and scaled with ScaleData; data dimension was reduced by Principal Component Analysis with RunPCA; Doublets were counted and removed with DoubletFinder^93^; the number of significant Principal Components (PC) was determined with the function JackStraw^94^; Uniform Manifold Approximation and Projection (UMAP) dimension reduction^95^ was performed with the function RunUMAP using the appropriate number of PCs. For the WT sample, MultiK^96^ was used to explore the optimal number of clusters; a nearest-neighbor graph was constructed with the function FindNeighbors; clustering was computed with the function FindClusters, using the parameters determined by MultiK; cluster markers were identified with the function FindMarkers. For the identification of clusters, orthology to *Arabidopsis thaliana* genes was defined as best or second best reciprocal Blast hit. A similar pipeline, up to clustering, was applied to the *star* and *wico* scRNA-Seq samples.

#### Data integration

To estimate the effect of protoplasting on WT scRNA-Seq clusters, genes differentially expressed after protoplasting with different log2FoldChange values (absolute log2FoldChange > 1, 1.5, 2 or 2.5) were removed from the scRNA-Seq dataset, to generate 4 datasets. These were integrated with the original WT dataset with Harmony^38^ (implemented in Seurat), and cell proportions per cluster were compared. To compare the WT, *star* and *wico* samples, the two biological replicates of WT scRNA-Seq were first integrated with Harmony; then the WT, *star* and *wico* samples (after computing the UMAP) were integrated together with Harmony, to ensure an unbiased representation of each genotype for cluster identification after integration.

#### Layer-specific gene differential expression

On the WT, *star* and *wico* datasets after integration, the 3 epidermal clusters and the 3 mesophyll clusters were merged together by redefining their identities, while the vasculature and replicating/dying cells were kept untouched. For the *star* sample only, the small cluster of limb epidermal cells (secondary L1-revertants) was removed from the epidermal cluster. Only genes with more than 50 total counts in the dataset were kept for further analysis. Layer-specific DEGs were computed with the function FindMarkers, and subsetted for an absolute log2FoldChange > 0.75 and layer-specificity > 10% (computed as pct.1 – pct.2, pct.1 being the percentage of cells from group 1 expressing the gene, and pct.2 from group 2). DEGs in the *star* epidermis and *wico* mesophyll, as compared to the corresponding tissue in WT, were computed similarly, and subsetted with an absolute log2FoldChange > 1, adjusted *p-*value < 0.01 and pct.1 - pct.2 >10 %. These cut-offs were chosen in an attempt to capture a meaningful number of genes, while being stringent for their layer-specificity of expression. Specifically, Seurat proposes a default cut-off of 0.25 for the absolute log2FoldChange, that we considered to be too permissive for our own data. The cut-off for layer-specificity of expression was chosen at 10%, as the lowest cut-off for which the numbers of epidermis- and mesophyll-enriched genes were statistically different in the WT petal. Venn diagrams were generated with the help of Interactivenn^97^.

### Chromatin immunoprecipitation and sequencing (ChIP-Seq)

#### ChIP assay and sequencing

Chromatin immunoprecipitation on WT, *star* and *wico* petals at stage 8 was performed as previously described^98,26^, using polyclonal antibodies directed against PhDEF devoid of its highly conserved DNA-binding domain. Initial enrichment tests by qPCR on two positive control genes (PhDEF and Pos2) and one negative control gene (Neg1) allowed to select the appropriate amount of sonicated chromatin, set at 25 µL, 100 µL and 100 µL for WT, *star* and *wico* respectively. Four replicate ChIP experiments were performed for each genotype, and enrichment was measured by ChIP-qPCR (Figure S10) to select the two best replicates to send for sequencing. The quality of IP and INPUT (WT chromatin) samples was assessed with Tapestation 4150 HS D5000 (Agilent). Libraries were prepared with MicroPlex Library Preparation Kit v3 (Diagenode) and around 1 ng of each library was PCR-amplified (12 cycles), following Diagenode’s recommendations. Libraries were analyzed TapeStation 4150 HS D5000 and quantified with Qubit 4.0 with the Qubit dsDNA HS Assay Kit (Thermofisher), then mixed at equimolar ratio for sequencing with an Illumina NextSeq 500 (paired-end, 2×76 bp, dual indexing). Read quality was assessed with FastQC^88^ and MultiQC^99^, and reads were trimmed with fastx-trimmer^100^ to remove the first 6 nucleotides on the 5’ end and the last nucleotide on the 3’ end of each read, resulting in 2×68 bp reads. Amplification bias was removed as previously described^101^. Reads were mapped on the *P. axillaris* genome (v1.6.2 superscaffolded by HiC^30,89,26^), which was the most recent version of the genome available at the time data analysis was initiated. Mapping was performed with Bowtie2^102^ (v2.4.2) with default parameters, and only reads with mapping quality above 20 were kept. The effective genome size was estimated at 9.99e8 with khmer^103^, with a kmer size of 68.

#### Peak detection and annotation

We explored different strategies for peak calling, but the standard MACS2 pipeline was not capturing PhDEF binding on its own promoter, which we used as our positive control, likely because the input sample was too noisy. Therefore, we explored different thresholds for peak calling with MACS2^104^ (False Discovery Rate = FDR) and for the Irreproducible Discovery Rate (IDR) ^40^ between IP replicates. Applying a FDR of 0.1 and an IDR of 0.1 to IP samples, and a FDR of 0.05 to INPUT samples, allowed to capture visually striking binding events in the IP and to remove spurious peaks in the INPUT in a satisfactory way. Peaks detected in the INPUT sample were removed from the reproducible IP peaks if they overlaped by at least 25 % reciprocally, or if INPUT peaks were included in IP peaks, which was computed with a custom script using a combination of bedtools^105^ (v2.26.0) commands (intersect, window, substract). Such customization of peak detection has been already applied to other transcription factors^106^. Annotation of genes close to peaks was performed with Python. In cases when a peak was assigned to several possible genes, the gene with the shortest distance to the peak was retained. Peak intersections were performed independently using bedtools intersect on (i) the complete peak sets from WT, *star*, and *wico*, and (ii) the subset of peaks located within 6 kb upstream of the transcription start site (TSS) or 6 kb downstream of the transcription termination site (TTS) of annotated genes, hereafter referred to as gene-associated peaks. Shared peaks were defined by a reciprocal overlap of at least 25%, in order to identify marked regions in one, two or three genotypes. The distance of 6 kb, which was retained for the rest of the analysis, was chosen based on the metaplots, and because it encompasses 80% of the ChIP peaks. Motifs under the peaks were explored with MEME-ChIP (v5.5.7)^107^ with a minimal size of 6 bp and a maximal size of 15 bp against the JASPAR 2024^108^ CORE non-redundant v2 database, and selected motifs (WT motifs from Fig. 3B) were used to predict binding sites on individual genomic sequences with FIMO^109^. Random genomic sequences, with the same size and size distribution as test files, were generated from the *P. axillaris* genome, for gene regions only or for the whole genome, to test for enrichment in TCP binding sites. Peaks and gene models were visualized with the Integrative Genome Viewer^110^. For heatmaps of read coverage, coverage files (bigwig files) were calculated from BAM files using bamCoverage from the package deepTools2^111^ (v3.5.4). Heatmaps were built using computeMatrix and plotHeatmap functions using a window of 3 kb beginning at the start of the peak.

### GO term database improvement and enrichment analysis

Protein sequences were predicted from the *P. axillaris* genome annotation transferred from the published version v1.6.2^84^ to the Hi-C superscaffolded one using GFFread^112^ and TransDecoder.LongOrfs^113^ (v5.5.0) with parameter “-m50”. A total of 56,351 predicted protein sequences were obtained. We then applied custom Perl scripts (refer to *Data availability* section) to select the most plausible CDS for each predicted mRNA, resulting in 31,626 predicted protein sequences. These sequences were then compared with UniProtKB protein database version 2024_06^114^. The complete Swiss-Prot database of curated proteins (containing 44,534 plant sequences and 571,609 non-plant sequences) and the plant subset of the noncurated database TrEMBL (containing 20,024,921 sequences) were used. The comparison was performed using BlastP^115^ program (AB-Blast v3.0 release 2020-03-17) with parameters “W=3 Q=7 R=2 matrix=BLOSUM80 B=200 V=200 E=1e-6 hitdist=60 hspsepqmax=30 hspsepsmax=30 sump postsw”. The BLAST output was filtered using custom Perl scripts to retain only those matches with a log10(*e*-value) no lower than 75% of the best log10(*e*-value). The GO annotations for proteins contained in the Uniprot KnowledgeBase were downloaded as a GPA file (v222, 2024-08-01) from the GOA database^116^ hosted at EBI. The Gene Ontology database^117^ was retrieved via the QuickGO web interface^118^ in October 2024; GO term relationships and UniprotKB GO annotations were subsequently loaded into a custom PostgreSQL database (v13.13) for easy querying. Custom Perl scripts were written to associate each predicted petunia protein sequence to the GO terms associated with all its matched proteins or with at least with five matched proteins. The comprehensive list of GO annotations for the entire Petunia proteome is accessible in the project dataset as a text file (see *Data availability* section), with the list of underlying Blast evidences. Enrichment analysis for test genes was performed using all genes expressed in the WT petal bulk RNA-Seq as background genes. A hypergeometric test (R version 4.2.2^92^) was applied to assess the significance of enrichment of each subset^119,120^. GO terms enriched (*p*-value < 0.01 with a hypergeometric test) and with at least 2 genes in the pathway were analyzed with REVIGO^121^ (Tiny mode) to reduce redundancy. Enriched GO terms for biological processes were sorted by their log10(*p*-value) and the 10 best terms were displayed in barplots generated with ggplot2^122^ in R.

## Data and code availability

All data and code are publicly available as of the date of publication. Bulk RNA-Seq raw fastq files and normalized counts after DESeq2 have been deposited at GEO (GSE336494). Single-cell RNA-seq raw fastq files, h5 matrices and rds images after integration have been deposited at GEO (GSE290697). ChIP-Seq fastq and BED files have been deposited at GEO (GSE308255). Scripts for bulk RNA-Seq analysis, GO database update and GO term enrichment, and scRNA-Seq analysis are available at gitbio.ens-lyon.fr/rdp/phdef_flower_layer. ChIP-Seq BAM and BAI files, ChIP-Seq and scRNA-Seq FastQC html reports, and functional annotation of the *Petunia axillaris* proteome are available at https://entrepot.recherche.data.gouv.fr/dataverse/PhDEF_Flower_layer.

