## Supplemental Data for "The petal identity gene PhDEF has a major binding and regulatory action in the epidermis"

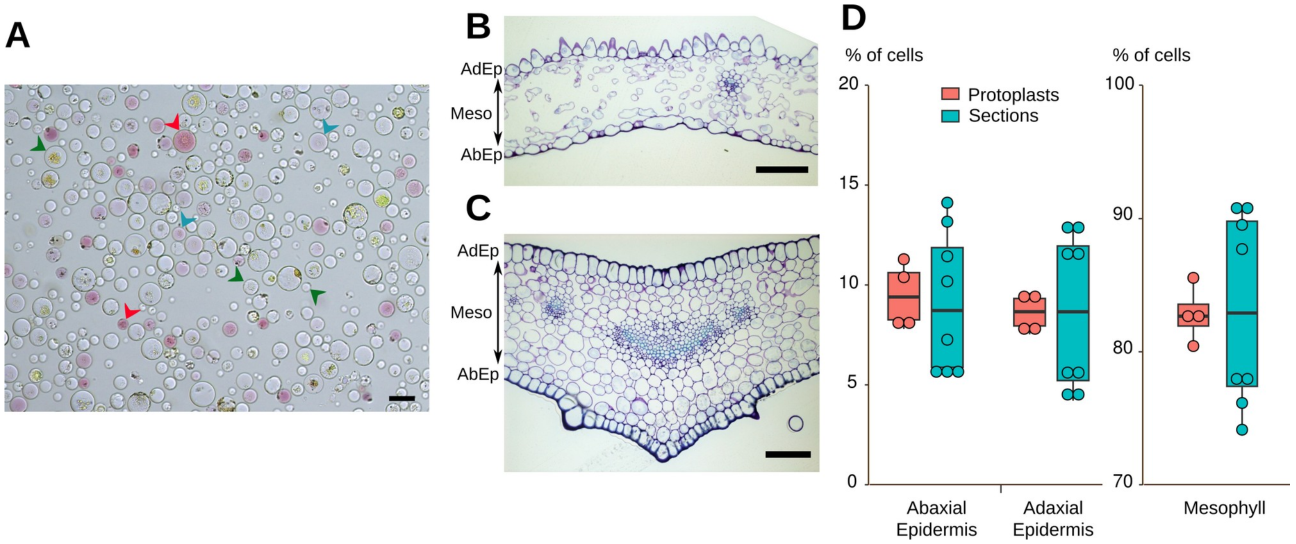


**Figure S1. Estimation of epidermal and mesophyll cell proportions from tissue sections or from petal protoplasting, Related to Figure 1.**

**(A)** Protoplasts isolated from WT mature petal. Strongly pigmented cells (like the ones pointed by the red arrowheads) are counted as cells from the adaxial epidermis, weakly pigmented cells (like the ones pointed by the blue arrowheads) are counted as cells from the abaxial epidermis, and the remaining cells (like the ones pointed by the green arrowheads) are counted as mesophyll cells. With this method, epidermal cells are under-estimated since tube epidermal cells are not pigmented. Scale bar = 50 µm. **(B,C)** Cross-sections from WT petal limb (B) and tube (C) after toluidine blue staining, allowing to count adaxial epidermis (AdEp), abaxial epidermis (AbEp) and mesophyll (Meso) cells. Scale bar = 100 µm. **(D)** Dotplot and boxplot of the percentage of cells from the adaxial epidermis, abaxial epidermis and mesophyll esimated from petal sections (tube and limb data merged) and from petal protoplast isolation. n = 4 pictures (each containing on average 834 cells for protoplasts, and 486 cells for tissue sections). Student's t-tests (two-sided) between protoplast and section cell numbers are non-significant in all cell layers.

**A**

| Sample | Forced number of cells | Mean reads per cell | Median UMI counts per cell | Median genes per cell | Number of reads | Valid barcodes | Sequencing saturation |
|---|---|---|---|---|---|---|---|
| **WT#1** | 4,000 | 277,689 | 3,270 | 1,394 | 1,110,754,946 | 93.2 % | 65.8 % |
| **WT#2** | 8,000 | 171,475 | 2,217 | 1,059 | 685,900,011 | 95.1 % | 84.3 % |
| ***star*#1** | 4,000 | 179,160 | 2,308 | 1,106 | 716,639,010 | 93.7 % | 59.3 % |
| ***wico*#1** | 4,000 | 271,723 | 1,678 | 794 | 1,086,891,925 | 93.7 % | 82.2 % |

| Sample | Reads mapped to genome | Reads confidently mapped to genome | Reads mapped confidently to intergenic regions | Reads mapped confidently to intronic regions | Reads mapped confidently to exonic regions | Reads mapped confidently to transcriptome | Fraction reads in cells | Total genes detected in scRNA-Seq | Total genes detected in bulk RNA-Seq |
|---|---|---|---|---|---|---|---|---|---|
| **WT#1** | 91.9 % | 35.1 % | 17.3 % | 2.8 % | 15.0 % | 16.6 % | 40.7 % | 22,215 | 24,361 |
| **WT#2** | 90.9 % | 69.2 % | 29.7 % | 6.6 % | 33.0 % | 38.4 % | 58.1 % | 21,749 | 24,361 |
| ***star*#1** | 89.0 % | 33.4 % | 16.1 % | 3.0 % | 14.3 % | 15.9 % | 60.1 % | 22,767 | 24,280 |
| ***wico*#1** | 82.1 % | 34.6 % | 15.2 % | 3.3 % | 16.2 % | 17.9 % | 66.0 % | 22,028 | 24,529 |

**B**

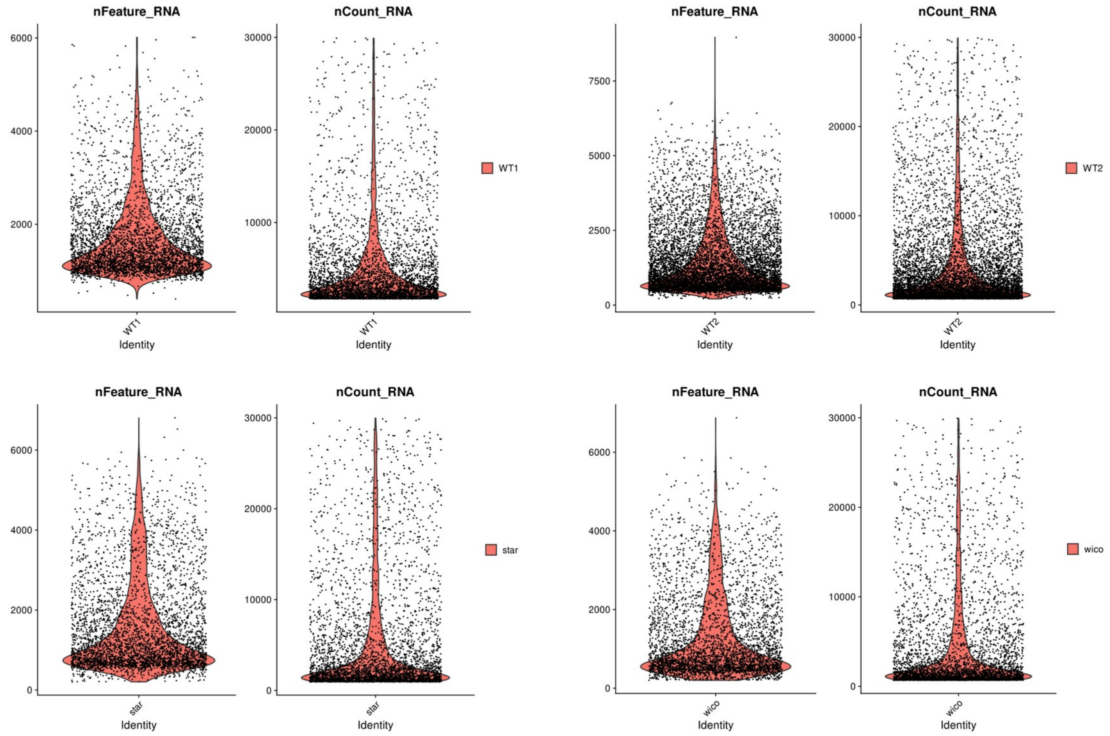


**Figure S2. Quality metrics of scRNA-Seq data. Related to Figure 1.**

**(A)** Quality and mapping metrics of the scRNA-Seq datasets produced in this study. Values are taken from the outputs of CellRanger. **(B)** Distribution of the number of unique features per cell (nFeature_RNA) and of the number of RNA counts per cell (nCount_RNA) for the two wild-type replicates (WT1 and WT2), the *star* and *wico* scRNA-Seq dataset.

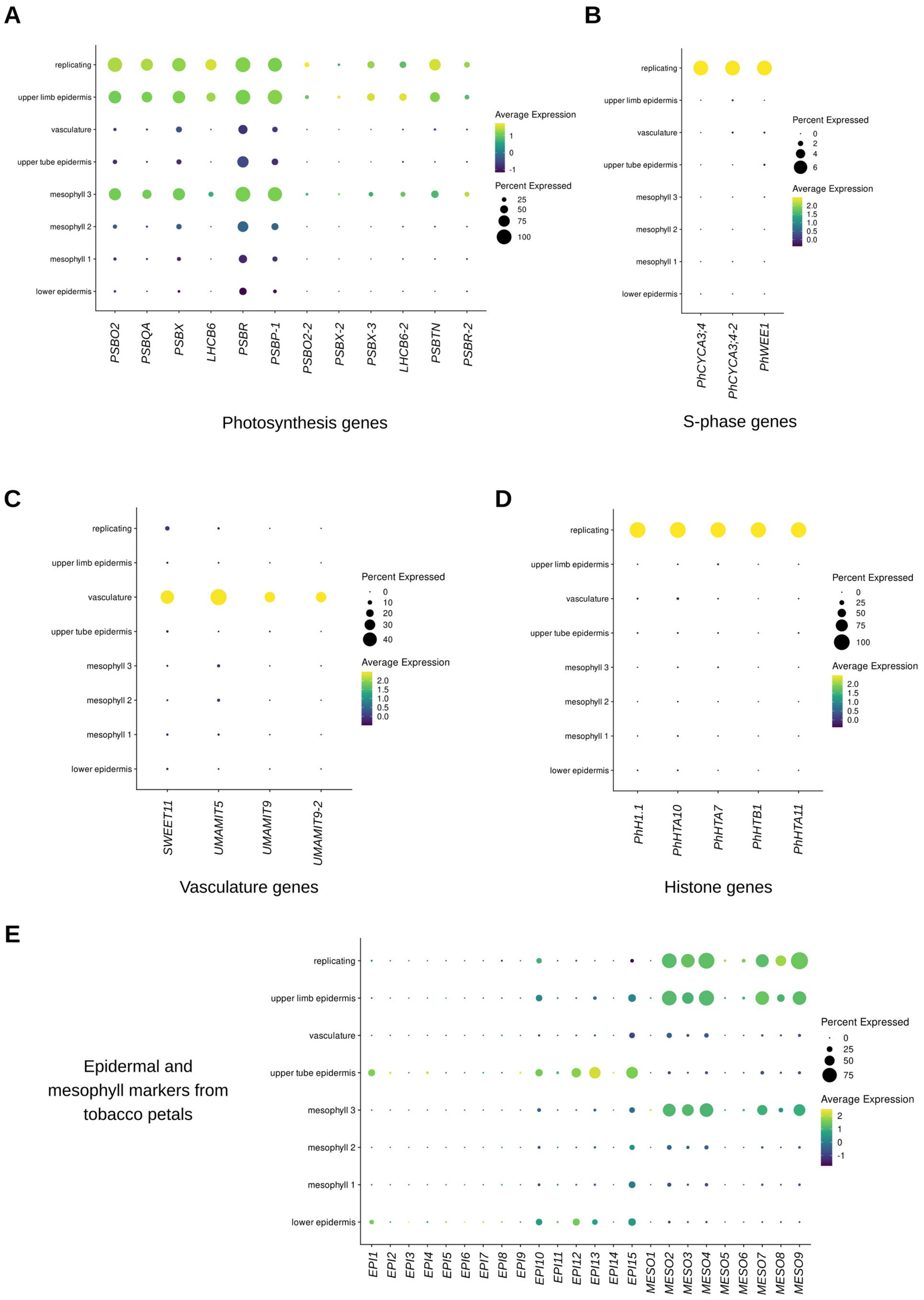
A
replicating
upper limb epidermis
vasculature
upper tube epidermis
mesophyll 3
mesophyll 2
mesophyll 1
lower epidermis
PSBO2
PSBQA
PSBX
LHCB6
PSBR
PSBP-1
PSBO2-2
PSBX-2
PSBX-3
LHCB6-2
PSBTN
PSBR-2
Average Expression
1
0
-1
Percent Expressed
25
50
75
100
Photosynthesis genes
B
PhCYCA3;4
PhCYCA3;4-2
PhWEE1
Percent Expressed
0
2
4
6
Average Expression
2.0
1.5
1.0
0.5
0.0
S-phase genes
C
SWEET11
UMAMIT5
UMAMIT9
UMAMIT9-2
Percent Expressed
0
10
20
30
40
Average Expression
2.0
1.5
1.0
0.5
0.0
Vasculature genes
D
PhH1.1
PhHTA10
PhHTA7
PhHTB1
PhHTA11
Percent Expressed
0
25
50
75
100
Average Expression
2.0
1.5
1.0
0.5
0.0
Histone genes
E
Epidermal and mesophyll markers from tobacco petals
EPI1
EPI2
EPI3
EPI4
EPI5
EPI6
EPI7
EPI8
EPI9
EPI10
EPI11
EPI12
EPI13
EPI14
EPI15
MESO1
MESO2
MESO3
MESO4
MESO5
MESO6
MESO7
MESO8
MESO9
Percent Expressed
0
25
50
75
Average Expression
2
1
0
-1

**Figure S3. Expression of selected marker genes in WT petal scRNA-Seq data. Related to Figure 1.**

**(A-E)** Dotplots of the expression of photosynthesis genes (Photosystem II subunits, **A**), S-phase genes (**B**), vasculature genes (**C**), histone genes (**D**), and epidermal (EPI) and mesophyll (MESO) markers from tobacco petals[34] (**E**). Gene identifiers can be found in Table S1.

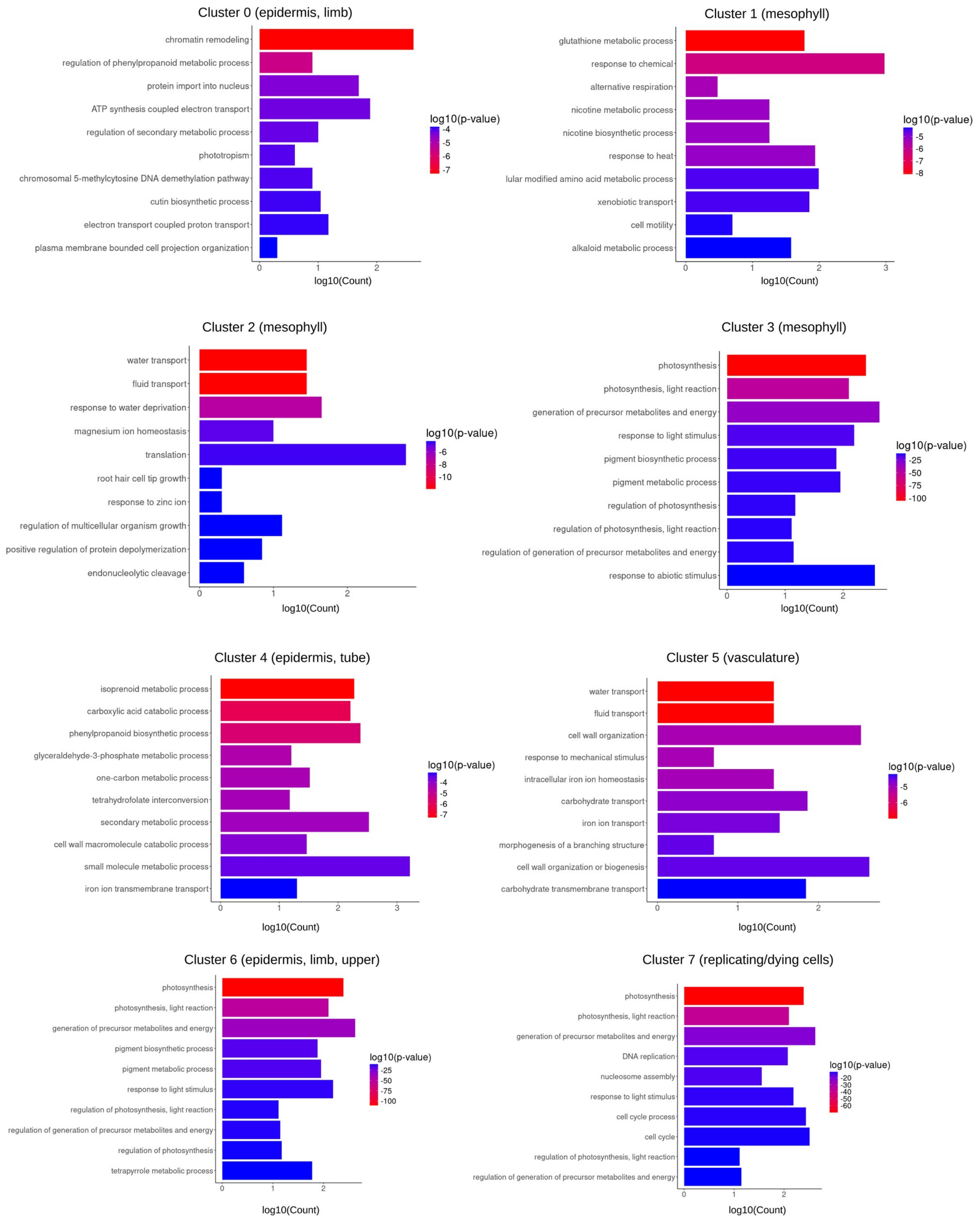


**Figure S4. Barplots of the ten best Gene Ontology (GO) terms enriched for the best marker genes of each cluster of WT petal cells. Related to Figure 1.**

Ten most enriched GO terms (Biological Process) for the best cluster markers (with *p*-adjusted value < 0.01 and log2FC > 1, as compared to all the other clusters) after redundancy reduction with REVIGO. The color indicates the significance of the enrichment (log10(*p*-value)) and the length of the bar indicates the number of genes associated to this GO term (log10(Count)). The full list of GO enriched terms is provided in Table S2.

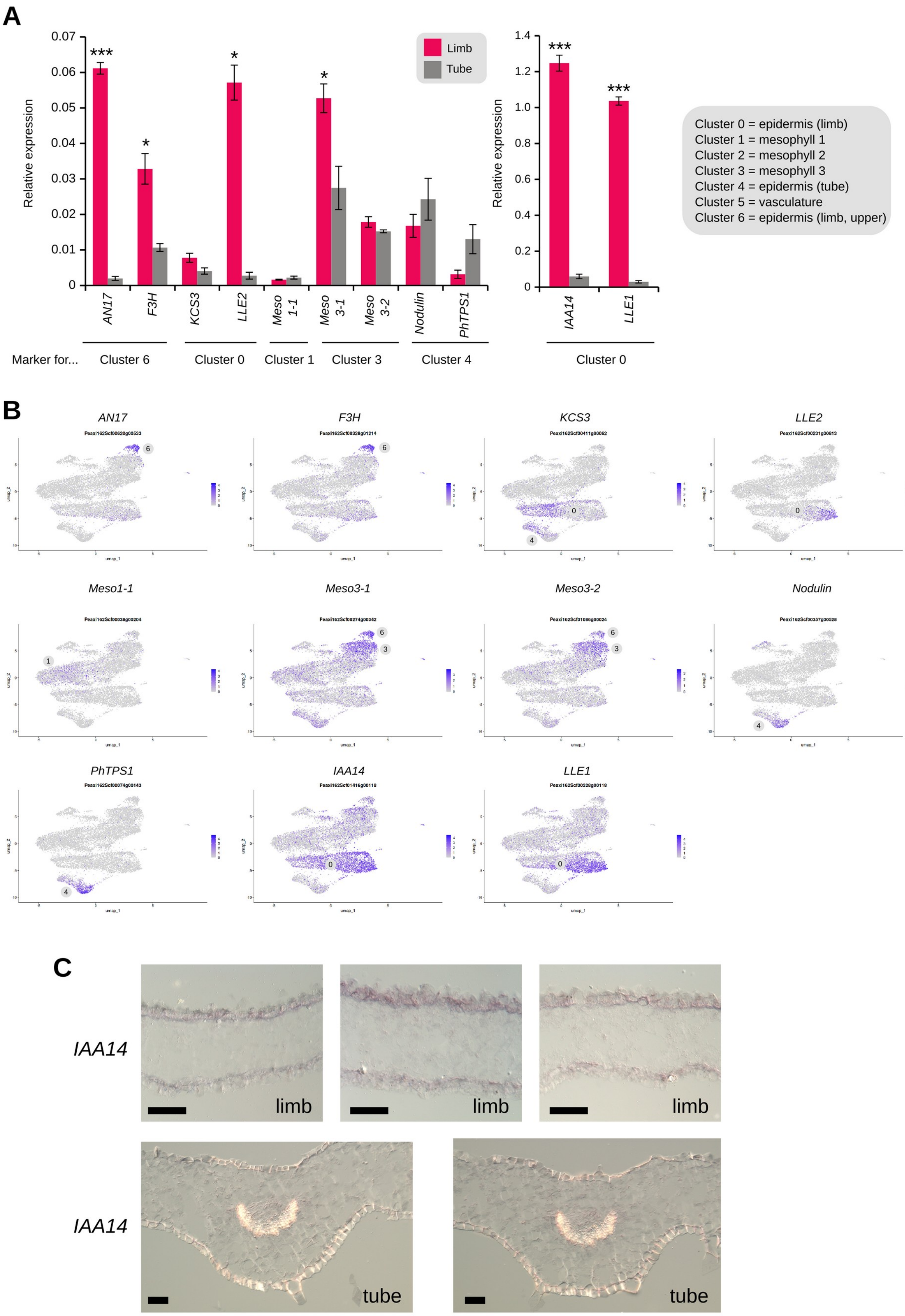
A
Relative expression
Limb
Tube
***
*
*
*
AN17
F3H
KCS3
LLE2
Meso 1-1
Meso 3-1
Meso 3-2
Nodulin
PhTPS1
IAA14
LLE1
Marker for...
Cluster 6
Cluster 0
Cluster 1
Cluster 3
Cluster 4
Cluster 0
Cluster 0 = epidermis (limb)
Cluster 1 = mesophyll 1
Cluster 2 = mesophyll 2
Cluster 3 = mesophyll 3
Cluster 4 = epidermis (tube)
Cluster 5 = vasculature
Cluster 6 = epidermis (limb, upper)
B
AN17
F3H
KCS3
LLE2
Meso1-1
Meso3-1
Meso3-2
Nodulin
PhTPS1
IAA14
LLE1
C
IAA14
limb
IAA14
tube

**Figure S5. Expression of cluster marker genes in limb and tube WT petal tissue and corresponding UMAP plots. Related to Figure 1.**

**(A)** Relative expression level of chosen cluster markers in limb and tube tissue from WT flowers at anthesis, measured by RT-qPCR. No suitable marker could be selected for mesophyll cluster 2. n = 3 biological replicates, mean ± s.e.m. Student's t-test two-sided or Wilcoxon rank sum exact test (for non-parametric data distribution for *KCS3* and *PhTPS1*), * $p < 0.05$, ** $p < 0.005$, *** $p < 0.001$.

**(B)** UMAP plots of gene expression in WT scRNA-Seq data for chosen cluster markers. Average expression is shown as a color scale set by default. For each plot, the cluster for which the gene is a marker is indicated. Gene identifiers are available in Table S2.

**(C)** *In situ* hybridization for the *IAA14* transcript, enriched in cluster 0, in limb (up) and tube (bottom) cross-sections. Sections are oriented with the upper epidermis at the top. A colorimetric signal is detected in the upper and lower epidermis in the limb, but not in the tube. Scale bar = 50 µm.

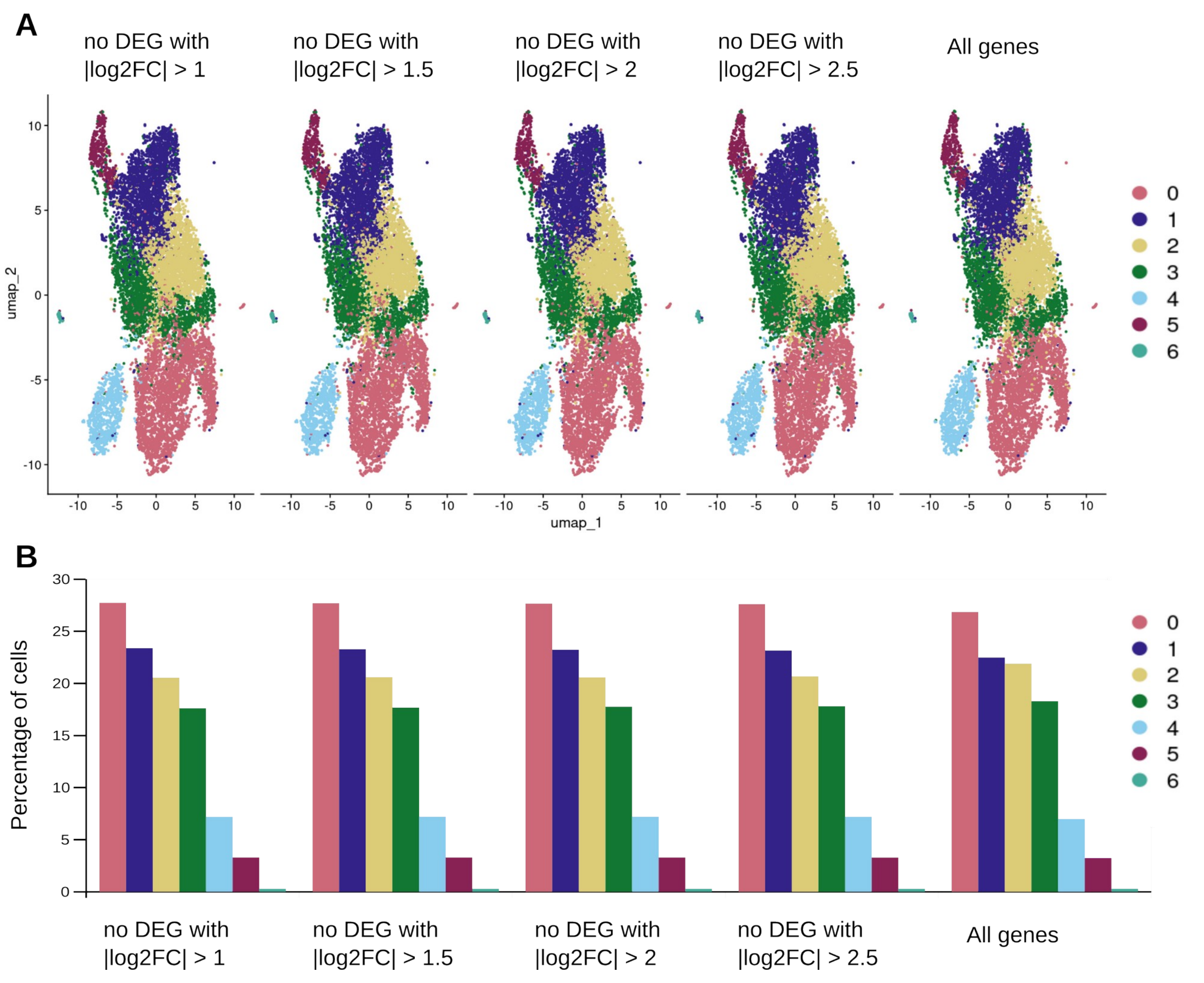


**Figure S6. Effect of removing DEGs by the protoplasting process on the scRNA-Seq UMAP and cell type proportions. Related to Figure 1.**

UMAP plots **(A)** and percentage of cells in each cluster **(B)** of WT petal scRNA-Seq data, removing genes differentially expressed (DEG) by the protoplasting process at different log2FoldChange cut-offs (|log2FC| = absolute value of log2FoldChange), or keeping all genes. The number of genes removed is the following : 9,985,7,646, 5,966 and 4,748 for |log2FC| values over 1.0, 1.5, 2.0 and 2.5, respectively. The color code for cell types is the same for the UMAP plots and the barplot, and corresponds to the cluster number. For this analysis, integration of datasets with Harmony slightly changes the UMAP shape and the identity of clusters, as compared to Figure 1C with the WT dataset with all genes, therefore clusters are not directly comparable.

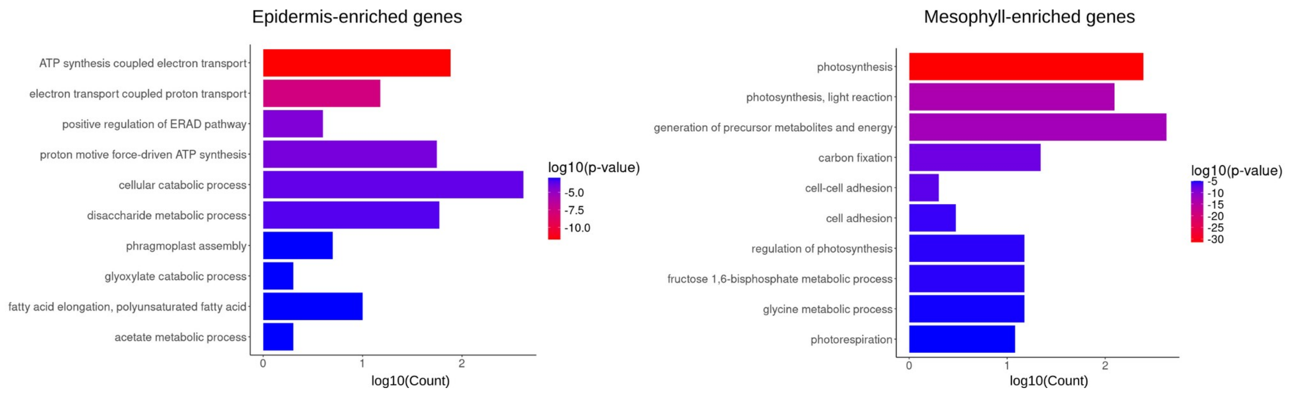


**Figure S7. Barplots of the ten best Gene Ontology (GO) terms enriched in the epidermis- and mesophyll-enriched genes. Related to Figure 1.**

Ten most enriched GO terms (Biological Process) for the epidermis- and mesophyll-enriched genes (selected with log2FC > 0.75, specificity of expression > 10 % and adjusted *p*-value < 0.01, as compared to the other layer) after redundancy reduction with REVIGO. The color indicates the significance of the enrichment (log10(*p*-value)) and the length of the bar indicates the number of genes associated to this GO term (log10(Count)). The full list of GO enriched terms are provided in Table S2.

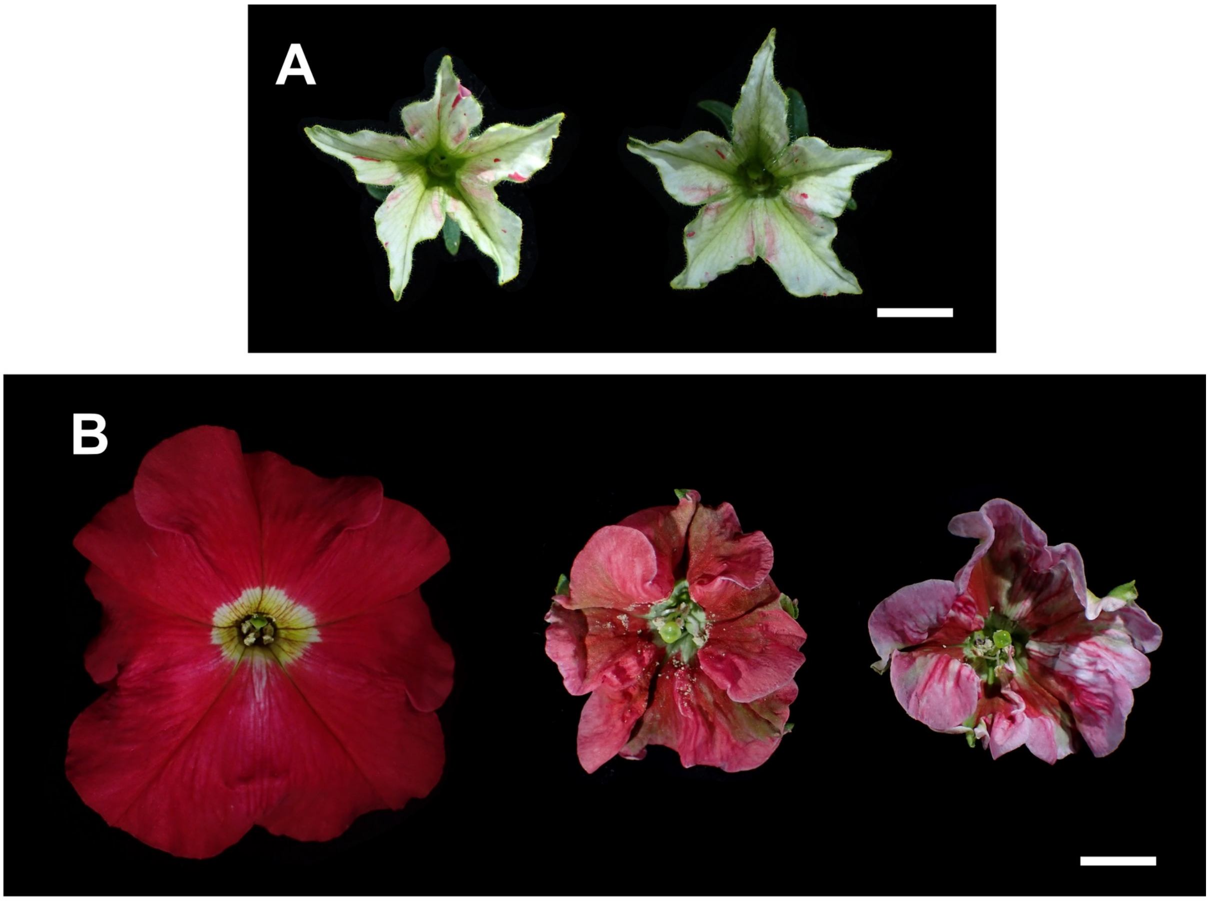


**Figure S8.** ***star*** **flowers frequently display pigmented sectors and** ***wico*** **flowers frequently show pigmentation defects. Related to Figure 2.**

**(A)** Small  pigmented L1-revertant sectors are visible in the petal limb of *star* flowers, due to *dTph1* excision in the epidermis, restoring *PhDEF* expression in clonal sectors. Scale bar = 1 cm.

**(B)** Two *wico* flowers with pigmentation defects (middle and right flowers), with a WT flower on the left for comparison. Scale bar = 1 cm.

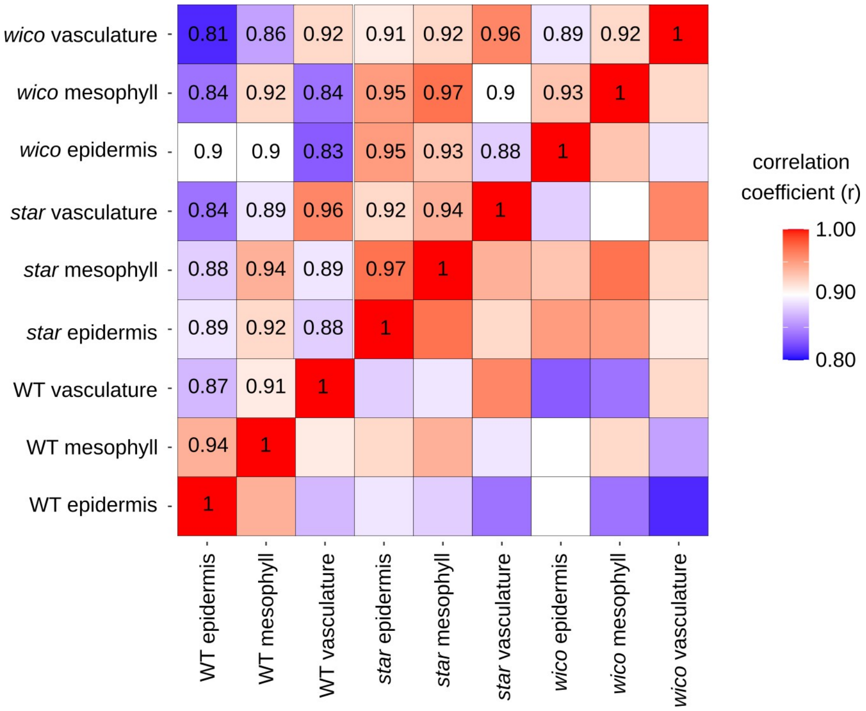


**Figure S9. Heatmap of correlation coefficients between pseudo-bulk transcriptomes from WT, *wico* and *star* epidermis, mesophyll and vasculature. Related to Figure 2.**

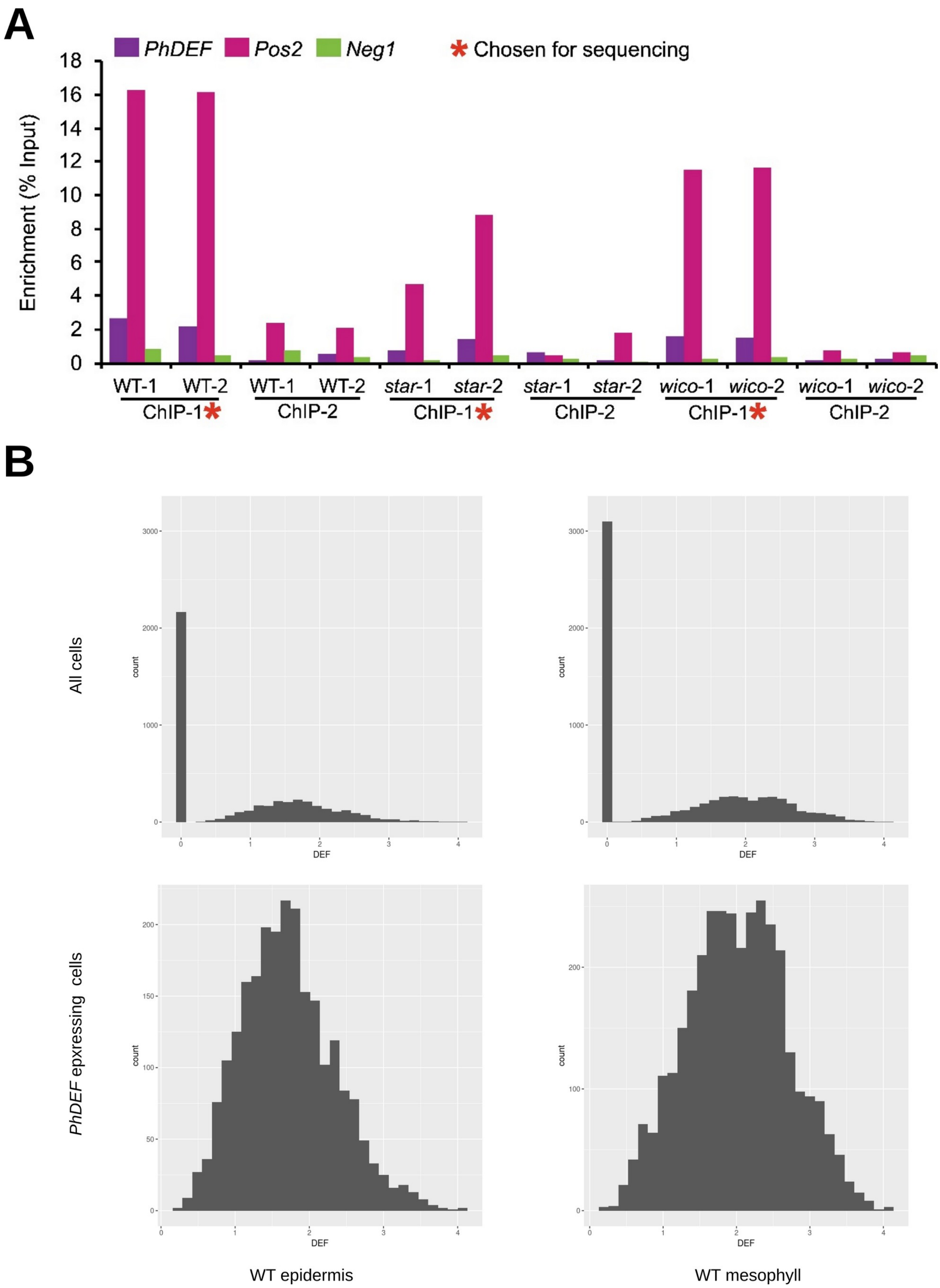


**Figure S10. Assessment of ChIP efficiency by qPCR, and analysis of variance of *PhDEF* expression levels in the WT epidermis and mesophyll.**

**(A)** Measure of ChIP enrichment in two ChIP assays (ChIP-1 and ChIP-2) for two replicates in each assay, for WT, *star* and *wico*. The enrichment is calculated as percentage of Input, measured by qPCR on two positive control binding sites (in the *PhDEF* promoter, and in the *Pos2* gene) and one negative control binding site (*Neg1*). The replicates from the first ChIP assay depicted a much better enrichment and were chosen for sequencing.

**(B)** Barplot of the number of cells (count, y-axis) expressing *PhDEF* at different levels (log-normalized read counts, x-axis) in the WT epidermis (left) and in the WT mesophyll (right), extracted from the WT scRNA-Seq data. The upper panels depict all cells, and the lower panels depict only cells that express *PhDEF*. The mean value of *PhDEF* expression is 0.89 in the epidermis and 1.06 in the mesophyll (1.71 and 2.01 after removing cells that do not express *PhDEF*, respectively, $p < 2.2 e^{-16}$, Wilocxon rank sum test, one-tailed). The coefficient of variation of *PhDEF* expression is 1.08 in the epidermis and 1.06 in the mesophyll (0.36 and and 0.34 after removing cells that do not express *PhDEF*, respectively, $p = 0.00106$ with Feltz and Miller' asymptotic test).

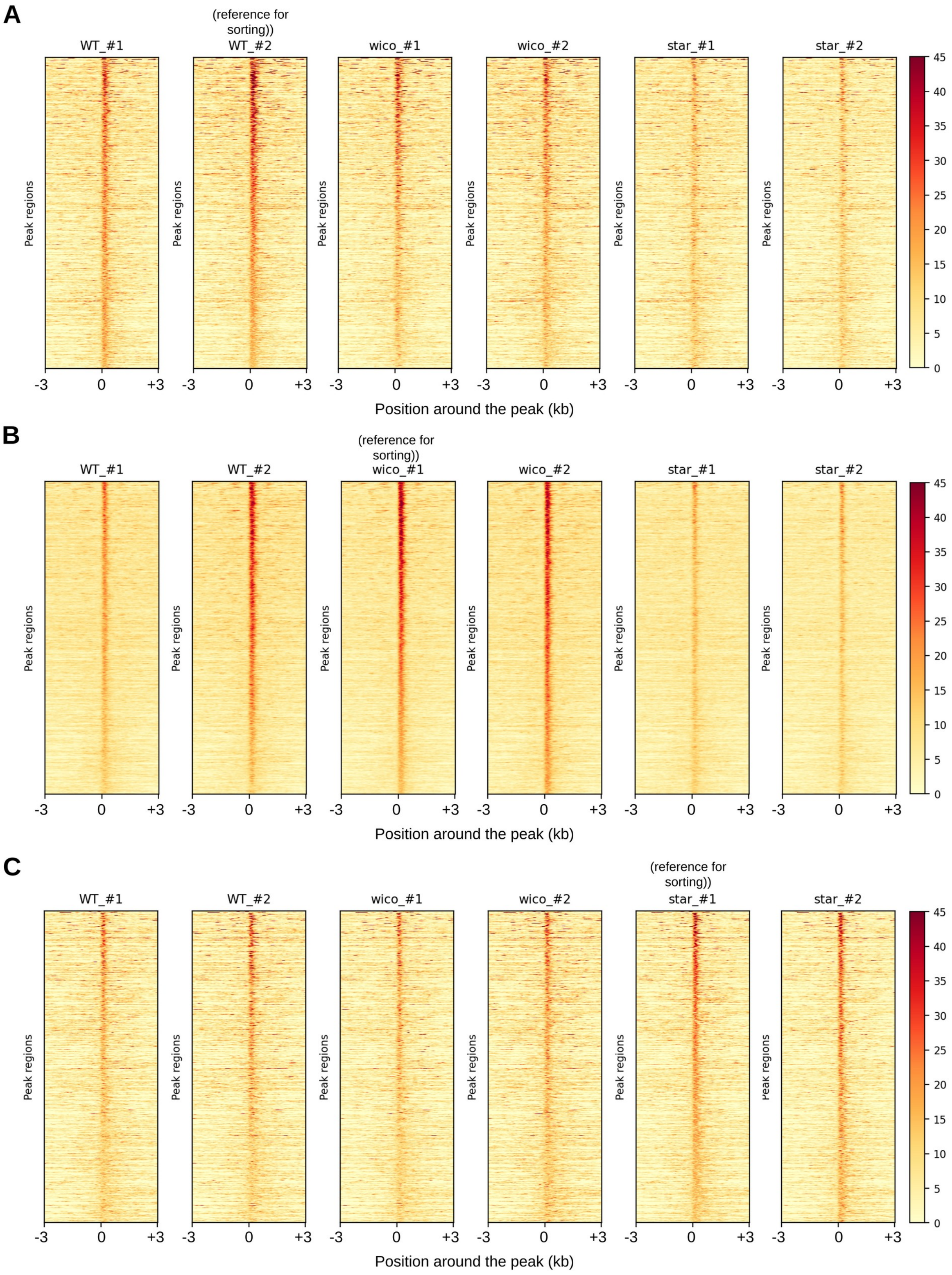


**Figure S11. Heatmap of the read coverage of peaks unique to a genotype. Related to Figure 3.** Heatmap of the read coverage of regions corresponding to peaks uniquely detected in WT (A), *wico* (B) and *star* (C) samples. The peaks are sorted according to the highest to lowest read coverage in WT #2 (A), *wico* #1 (B) and *star* #1 (C), so that the same regions of the genome are in the same horizontal line in all librairies. The read coverage is color-coded, and position 0 represents the start of the peak.

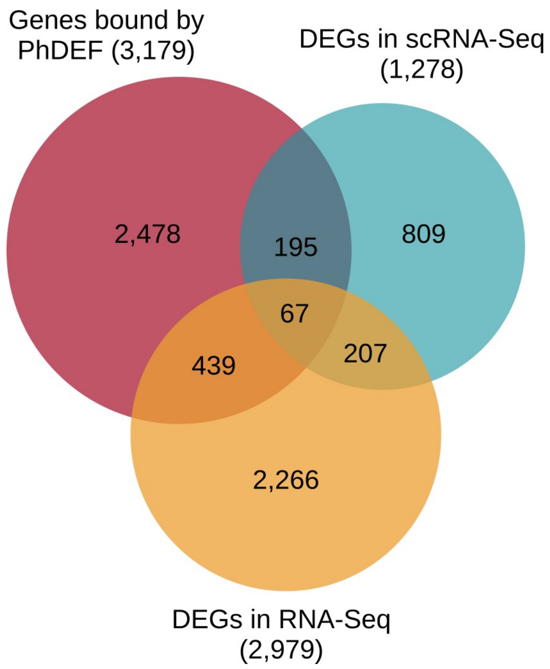


**Figure S12. Venn diagram depicting the intersection of the ChIP-Seq, bulk RNA-Seq and scRNA-Seq datasets.**

Genes bound by PhDEF are all genes bound by PhDEF in ChIP-Seq in WT, *wico* and/or *star* at stage 8. DEGs in RNA-Seq are DEGs in *star* (stage 8) and/or *wico* (stage 8) as compared to WT (stage 8). DEGs in scRNA-Seq are DEGs in *star* epidermis vs. WT epidermis and/or in *wico* mesophyll vs. WT mesophyll in mature petals. The overlap between genes bound by PhDEF and DEGs in RNA-Seq is significant (506 genes, $p = 4.73e^{-13}$, hypergeometric test), the overlap between genes bound by PhDEF and DEGs in scRNA-Seq is significant (262 genes, $p = 1.37e^{-04}$), and the overlap between DEGs in RNA-Seq and DEGs in scRNA-Seq is significant (274 genes, $p = 2.52e^{-17}$).

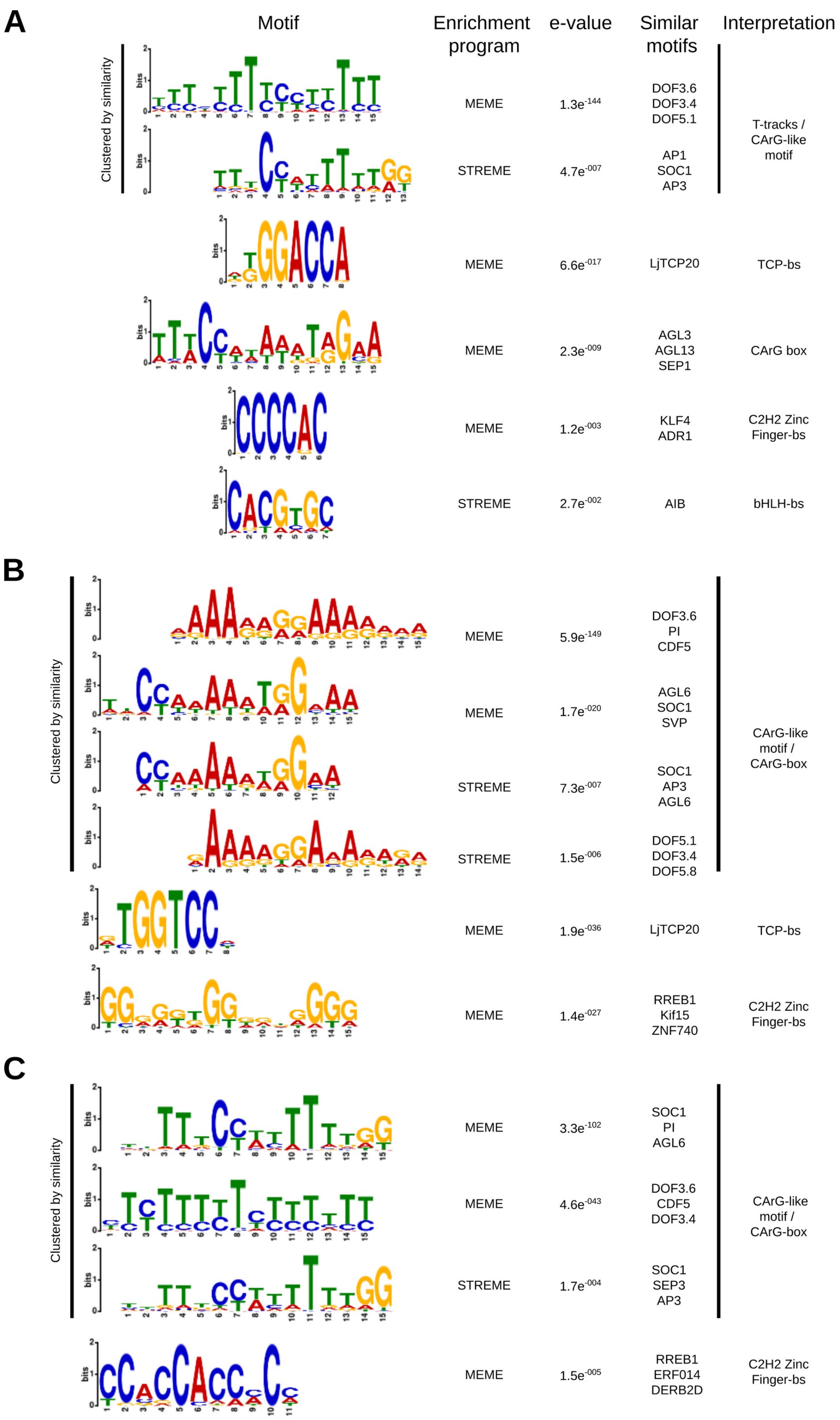
A
Motif
Enrichment program
e-value
Similar motifs
Interpretation
Clustered by similarity
MEME
1.3e-144
DOF3.6
DOF3.4
DOF5.1
STREME
4.7e-007
AP1
SOC1
AP3
T-tracks / CArG-like motif
MEME
6.6e-017
LjTCP20
TCP-bs
MEME
2.3e-009
AGL3
AGL13
SEP1
CArG box
MEME
1.2e-003
KLF4
ADR1
C2H2 Zinc Finger-bs
STREME
2.7e-002
AIB
bHLH-bs
B
Clustered by similarity
MEME
5.9e-149
DOF3.6
PI
CDF5
MEME
1.7e-020
AGL6
SOC1
SVP
STREME
7.3e-007
SOC1
AP3
AGL6
STREME
1.5e-006
DOF5.1
DOF3.4
DOF5.8
CArG-like motif / CArG-box
MEME
1.9e-036
LjTCP20
TCP-bs
MEME
1.4e-027
RREB1
Kif15
ZNF740
C2H2 Zinc Finger-bs
C
Clustered by similarity
MEME
3.3e-102
SOC1
PI
AGL6
MEME
4.6e-043
DOF3.6
CDF5
DOF3.4
STREME
1.7e-004
SOC1
SEP3
AP3
CArG-like motif / CArG-box
MEME
1.5e-005
RREB1
ERF014
DERB2D
C2H2 Zinc Finger-bs

**Figure S13. Full output of MEME-chip with motifs significantly enriched below ChIP peaks in WT, *wico* and *star*. Related to Figure 3.**

MEME-chip output for WT (A), *wico* (B) and *star* (C) ChIP-Seq peaks associated with genes. The column « similar motifs » corresponds to motifs automatically proposed by MEME-chip based on similarity with the JASPAR 2024 database, while the column « Interpretation » is our own interpretation of the motif based on the family of transcription factors found to be similar.

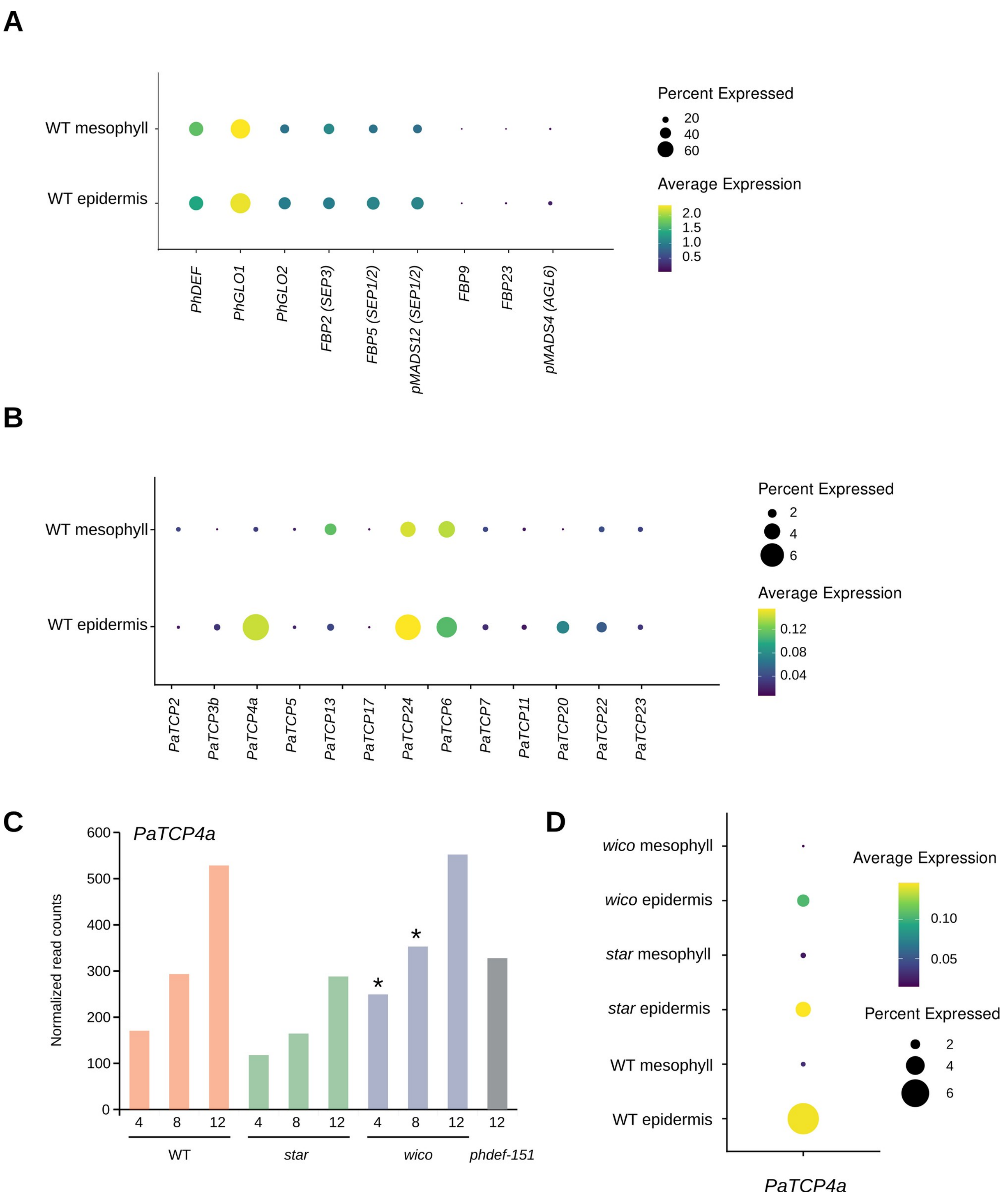


**Figure S14. Expression level of B- and E-class MADS-box genes and *TCP* genes in scRNA-Seq and bulk RNA-Seq. Related to Figure 3.**

**(A)** Dotplot of the expression of B- and E-class MADS-box genes in the WT epidermis and mesophyll. For E-class genes, the name of the *Arabidopsis thaliana* ortholog is indicated between parentheses, from[22]. **(B)** Dotplot of the expression of *TCP* genes in the WT epidermis and mesophyll. The name and identifiers of *Petunia axillaris TCP* genes originate from[50]. The *TCP* genes that are not displayed here were not found to be expressed. **(C)** Expression level of *PaTCP4a* in petals from WT, *star* and *wico*, from bulk RNA-Seq at different stages (4, 8 and 12). Stars indicate a significantly higher expression in *wic*o than in *star*, as computed by DESeq2 (log2FC > 1, *p*-adj < 0.01). **(D)** Dotplot of *PaTCP4a* expression in WT, *star* and *wico* epidermis and mesophyll, from scRNA-Seq data at mature stage.

| | *Petunia axillaris* | | *Arabidopsis thaliana* |
|---|---|---|---|
| **Marker for…** | **Gene ID** | **Gene Name in Figure 1D and S3** | **Gene ID** |
| **Epidermis** | Peaxi162Scf00411g00062 | *KCS3* (this study) | AT1G07720 |
| | Peaxi162Scf00814g00227 | *KCS21* (this study) | AT5G49070 |
| | Peaxi162Scf00922g00110 | *PFI* (this study) | AT2G26250 |
| | Peaxi162Scf00788g00213 | *PFI-2* (this study) | AT2G26250 |
| | Peaxi162Scf00021g00922 | *HDG1* (this study) | AT3G61150 |
| | Peaxi162Scf00262g00121 | *HDG12* (this study) | AT1G17920 |
| | Peaxi162Scf00950g00058 | *HDG12-2* (this study) | AT1G17920 |
| | Peaxi162Scf01416g00118 | *IAA14* (this study) | AT4G14550 |
| | Peaxi162Scf00328g00118 | *LLE1* (this study) | |
| | Peaxi162Scf00231g00813 | *LLE2* (this study) | AT2G18890 |
| **Tube upper epidermis** | Peaxi162Scf00074g00143 | *PhTPS1* | AT1G48820 |
| | Peaxi162Scf00357g00528 | *Nodulin* (this study) | |
| **Vasculature** | Peaxi162Scf00647g00518 | *SWEET11* (this study) | AT3G48740 |
| | Peaxi162Scf00656g00026 | *UMAMIT5* (this study) | AT1G75500 |
| | Peaxi162Scf00601g00047 | *UMAMIT9* (this study) | AT5G07050 |
| | Peaxi162Scf00003g05227 | *UMAMIT9-2* (this study) | AT5G07050 |
| **Upper limb epidermis** | Peaxi162Scf00620g00533 | *ANS/AN17* | AT4G22880 |
| | Peaxi162Scf00328g01214 | *F3H* | AT3G51240 |
| | Peaxi162Scf00536g00092 | *CHSj* | AT5G13930 |
| | Peaxi162Scf00518g00430 | *MT* | AT1G67980 |
| | Peaxi162Scf00713g00038 | *AN9* | AT5G17220 |
| | Peaxi162Scf00378g00113 | *5GT* | AT4G14090 |
| | Peaxi162Scf00858g00215 | *PALa* | AT3G10340 |
| | Peaxi162Scf00487g00064 | *RT* | AT1G50580 |
| | Peaxi162Scf03779g00019 | *MYB27* | AT2G16720 |
| | Peaxi162Scf00578g00007 | *AN4* | AT1G56650 |
| | Peaxi162Scf00163g00081 | *3GT* | AT5G17050 |
| | Peaxi162Scf00521g00814 | *MYBx* | AT1G56650 |
| | Peaxi162Scf00488g00074 | *PALb* | AT3G10340 |
| **Replicating/Dying cells** | Peaxi162Scf00813g00127 | *H1.1* (this study) | AT1G06760 |
| | Peaxi162Scf00037g00927 | *HTA10* (this study) | AT1G51060 |
| | Peaxi162Scf00071g00634 | *HTA7* (this study) | AT5G27670 |
| | Peaxi162Scf00114g00053 | *HTB1* (this study) | AT1G07790 |
| | Peaxi162Scf01039g00231 | *HTA11* (this study) | AT3G54560 |
| | Peaxi162Scf00160g00031 | *CYCA3;4* (this study) | AT1G47230 |
| | Peaxi162Scf00848g00318 | *CYCA3;4-2* (this study) | AT1G47230 |
| | Peaxi162Scf00067g00235 | *WEE1* (this study) | AT1G02970 |
| **Mesophyll** | Peaxi162Scf00038g00204 | *Meso1-1* (this study) | |
| | Peaxi162Scf00274g00342 | *Meso3-1* (this study) | AT5G61410 |
| | Peaxi162Scf01086g00024 | *Meso3-2* (this study) | AT5G54600 |
| **Photosynthesis (Photosystem II subunits)** | Peaxi162Scf00019g00219 | *PSBO2* (this study) | AT3G50820 |
| | Peaxi162Scf00082g01216 | *PSBQA* (this study) | AT4G21280 |
| | Peaxi162Scf00101g00142 | *PSBX* (this study) | AT2G06520 |
| | Peaxi162Scf00152g01119 | *LHCB6* (this study) | AT1G15820 |
| | Peaxi162Scf00232g01129 | *PSBR* (this study) | AT1G79040 |
| | Peaxi162Scf00359g01218 | *PSBP-1* (this study) | AT1G06680 |
| | Peaxi162Scf00589g00517 | *PSBO2-2* (this study) | AT3G50820 |
| | Peaxi162Scf00724g00052 | *PSBX-2* (this study) | AT2G06520 |
| | Peaxi162Scf00928g00002 | *PSBX-3* (this study) | AT2G06520 |
| | Peaxi162Scf00001g00212 | *LHCB6-2* (this study) | AT1G15820 |
| | Peaxi162Scf00004g00263 | *PSBTN* (this study) | AT3G21055 |
| | Peaxi162Scf00006g00149 | *PSBR-2* (this study) | AT1G79040 |

| | **Gene ID** | **Gene Name in Figure S3** | **Tobacco ID** |
|---|---|---|---|
| **Tobacco petal epidermis** | Peaxi162Scf00595g00212 | *EPI1* (this study) | A4A49_30556 |
| | Peaxi162Scf00467g00920 | *EPI2* (this study) | A4A49_08035 |
| | Peaxi162Scf00755g00009 | *EPI3* (this study) | A4A49_66108 |
| | Peaxi162Scf00008g02420 | *EPI4* (this study) | A4A49_30297 |
| | Peaxi162Scf00049g02410 | *EPI5* (this study) | A4A49_28379 |
| | Peaxi162Scf00049g02514 | *EPI6* (this study) | A4A49_28379 |
| | Peaxi162Scf00141g00059 | *EPI7* (this study) | A4A49_30297 |
| | Peaxi162Scf00251g00123 | *EPI8* (this study) | A4A49_28379 |
| | Peaxi162Scf00394g00426 | *EPI9* (this study) | A4A49_66108 |
| | Peaxi162Scf00544g00513 | *EPI10* (this study) | A4A49_66108 |
| | Peaxi162Scf00786g00434 | *EPI11* (this study) | A4A49_22795 |
| | Peaxi162Scf00788g00213 | *EPI12* (this study) | A4A49_29650 |
| | Peaxi162Scf00922g00110 | *EPI13* (this study) | A4A49_29650 |
| | Peaxi162Scf00977g00015 | *EPI14* (this study) | A4A49_08850 |
| | Peaxi162Scf01343g00117 | *EPI15* (this study) | A4A49_28379 |
| **Tobacco petal mesophyll** | Peaxi162Scf00000g00601 | *MESO1* (this study) | A4A49_09113 |
| | Peaxi162Scf00035g02214 | *MESO2* (this study) | A4A49_09113 |
| | Peaxi162Scf00051g00910 | *MESO3* (this study) | A4A49_26955 |
| | Peaxi162Scf00045g02242 | *MESO4* (this study) | A4A49_02826 |
| | Peaxi162Scf01099g00035 | *MESO5* (this study) | A4A49_05181 |
| | Peaxi162Scf01311g00015 | *MESO6* (this study) | A4A49_05181 |
| | Peaxi162Scf01311g00019 | *MESO7* (this study) | A4A49_05181 |
| | Peaxi162Scf01311g00120 | *MESO8* (this study) | A4A49_05181 |
| | Peaxi162Scf01311g00122 | *MESO9* (this study) | A4A49_05181 |

**Table S1. Main cluster marker genes used to identify scRNA-Seq clusters. Related to Figure 1.** First sub-table: list of *Petunia axillaris* and *Arabidopsis thaliana* gene identifiers (ID) and gene names enriched in WT scRNA-Seq clusters, and used to assign cell type identity to clusters. Second sub-table: list of *Petunia axillaris* and *Nicotiana attenuata* (tobacco) gene identifiers (ID) and gene names, from epidermis-enriched and mesophyll-enriched genes in the tobacco petal[34]. Gene orthology was inferred by best or second best reciprocal Blast hit, as decribed in the Methods section. "This study" indicates that the gene name had not been assigned previously in petunia.

**Table S2 (available as separate file). Gene Ontology terms enriched in the best cluster markers from WT petal cells, in the layer-enriched DEGs and in the layer-specific DEGs. Related to Figure 1 and Figure 2.**

List of enriched GO terms for the best cluster markers (with adjusted *p*-value < 0.01 and log2FC > 1, as compared to all the other clusters) of WT petal cells, in the layer-enriched DEGs (with adjusted *p*-value < 0.01, log2FC > 0.75 and specificity of expression > 10 %, as compared to the other layer), and in the layer-specific DEGs (differentially expressed in the *star* epidermis, in the *wico* mesophyll, or in both for the common DEGs). Each sheet contains the GO term and its description, the count and fraction of genes associated to this term in the subset and in the control (all genes expressed in the petal) and the enrichment ratio and *p*-value for enrichment. The terms highlighted in yellow are the ones retained after reduction of redundancy with REVIGO, and displayed on Figures 2H, S4 and S7.

**Table S3 (available as separate file). List of differentially expressed genes after WT petal protoplasting. Related to Figure 1.** Total RNA was sequenced from WT petal tissue, with or without protoplasting. Differential expression was computed with DESeq2. log2FoldChange = fold change of a given gene in protoplasted vs. crude WT petal tissue, expressed in log2 value ; lfcSE = standard error of log2FoldChange ; padj = *p*-value adjusted for multiple testing ; GOterms = GO terms associated with a given gene; Annotation = automatic annotation from[40]; Manual Annotation = manual annotation of pigmentation-related genes and transcription factors of interest for petal identity and development.

**Table S4 (available as separate file). List of layer-enriched genes and genes with layer-specific differential expression in *phdef* mutant cell layers. Related to Figures 1 and 2.**

The content of the table is explained in the first sheet. The second sheet contains the list of layer-enriched genes (as determined with the WT scRNA-Seq data), defined as having an absolute log2(FoldChange) > 0.75 and a specificity of expression > 10 %; a positive log2(FoldChange) indicates that the gene is enriched in the epidermis, and a negative one that the gene is enriched in the mesophyll. The third, fourth and fifth sheets contain respectively the lists of genes that are differentially expressed in the *star* epidermis, the *wico* mesophyll, and both. These represent potential direct and indirect targets of PhDEF in individual cell layers.

**Table S5 (available as separate file). ChIP metrics and position of all ChIP peaks detected in WT, *star* and *wico* ChIP-Seq. Related to Figure 3.**

The first sheet contains ChIP mapping metrics for each library: raw number of reads, number of unique reads, percentage of duplicated reads, percentage of mapping, number of reads after mapping quality filtering, number of mapped reads in IDR peaks, and FriP score (fraction of reads in peaks). The following sheets contain the position of the peak in the genome (Chromosome, start, end) and the peak identifier (PEAK_ID) for all genotypes and their intersections.

| **Number of peaks (%)** | **WT** | ***wico*** | ***star*** |
|---|---|---|---|
| promoter (6kb) | 1,080 (34.5%)*** | 1,550 (36.2%)*** | 542 (24.5%)*** |
| terminator (6kb) | 630 (20.1%)*** | 844 (19.7%)*** | 410 (18.5%)*** |
| exon | 117 (3.7%) | 126 (2.9%) | 54 (2.4%) |
| intron | 190 (6.1%) | 251 (5.9%) | 130 (5.9%) |
| intergenic | 895 (28.6%) | 1,223 (28.6%) | 824 (37.2%) |
| N/A | 222 (7.1%) | 282 (6.6%) | 256 (11.6%) |
| Total | 3,134 | 4,276 | 2,216 |

| **Number of genes (%)** | **WT** | ***wico*** | ***star*** |
|---|---|---|---|
| with a single ChIP peak | 1,692 (91.5%) | 2,227 (89.5%) | 1,017 (94.5%) |
| with 2 ChIP peaks | 150 (8.1%) | 243 (9.8%) | 58 (5.4%) |
| with 3 ChIP peaks | 7 (0.4%) | 18 (0.7%) | 1 (0.1%) |
| with 4 ChIP peaks | 1 (0.05%) | 1 (0.04%) | 0 (0.0%) |
| Total | 1,850 | 2,489 | 1,076 |

**Table S6. Number of peaks and genes from the WT, *star and wico* ChIP-Seq experiment. Related to Figure 3.**

Upper table : Number and percentage of peaks associated with genes in their promoter (6 kb upstream of the TSS), terminator (6 kb downstream of the TTS), exon or intron, in intergenic regions or unannotated regions of the genome (N/A), from the WT, *star* and *wico* ChIP-Seq experiment. Peak enrichment was computed with a hypergeometric test, *** p< 0.001.

Lower table : Number and percentage of genes with 1 to 4 ChIP peaks detected, from the WT, *star* and *wic*o ChIP-Seq experiment.

**Table S7 (available as separate file). Position of all gene-associated ChIP peaks detected in WT, *star* and *wico* ChIP-Seq, and their intersections. Related to Figure 3.**

Each table contains the identifier of the peak (Domain), its position in the genome (Chromosome, start, end), the associated gene (as defined with the shortest distance to the peak), the position of the gene (start, end), the shortest distance of the peak to the gene, and the annotation of the peak position related to the gene (promoter, terminator, exon or intron). The first 3 sheets contain the gene-associated peaks found in WT, *wico* and *star* ChIP-Seq, and the following sheets contain the list of peaks detected in a single genotype or detected in the different intersections (WT-*wico*, WT-*star*, *wico*-*star*, WT-*wico*-*star*).

**Table S8 (available as separate file). Analysis of enrichment of TCP binding sites. Related to Figure 3.**

TCP binding sites were predicted with Fimo under epidermal-specific PhDEF binding peaks (Epi), mesophyll-specific (Meso) or shared peaks. Binding sites were either predicted for all peaks, or for gene-associated peaks only. Control sequences were either random DNA sequences artificially generated, or random sequences taken under genes from the *P. axillaris* genome (to compare with gene-associated peaks) or taken under random regions of the genome excluding telomeres and centromeres (to compare with all peaks. These control sequences were design to have the same distribution of lengths and total size than PhDEF binding peaks. A Fisher's exact test was applied to compare observed and expected binding sites.

| Gene | Primer name | Orientation | Sequence | Purpose |
|---|---|---|---|---|
| ***AN17*** | **MLY2994** | fw | CCAATGGCAATGTCCAAGGCTACG | RT-qPCR |
| | **MLY2995** | rv | CTTGTTGCTGGAGTGTAGTCAGTAGG | |
| ***F3H*** | **MLY2996** | fw | CCATCTACAGGGTGAAGTGGTCCAAGA | |
| | **MLY2997** | rv | GACATCCAATAACTTGCAAGCTAACTCCA | |
| ***Nodulin*** | **MLY2998** | fw | GGAGTAGTTGTAGGGTCAATCTTGACTGT | |
| | **MLY2999** | rv | GGATCCTGAGGTGATTGCAATTCCAAAT | |
| ***PhTPS1*** | **MLY3000** | fw | GGGCATGAAGTTGAACAAGAAAGAGGAC | |
| | **MLY3001** | rv | GGCGCTTCAGTTGCTCCAAGG | |
| ***LLE1*** | **MLY3002** | fw | CGCAATCTCTCGGAGGAGGACTAC | |
| | **MLY3003** | rv | GCTGGCTGGTTTAGATAAGCATATTTGC | |
| ***LLE2*** | **MLY3004** | fw | CAAATCAGAAAATGTGGTTGGGAGAGGAG | |
| | **MLY3005** | rv | CACATTAGGATGACAAACGTGACCAAGTG | |
| ***IAA14*** | **MLY3006** | fw | GAGGATTTTCTGAGACTGTTGATTTGAAACTC | |
| | **MLY3007** | rv | CTTGTGCCTTGGCTGGTGGCTT | |
| ***KCS3*** | **MLY3008** | fw | GAATGGTATTTGATGGTCGTGAATCATGTCC | |
| | **MLY3009** | rv | GCATGGATATATTCACCACGAGGACATC | |
| ***Meso1-1*** | **MLY3014** | fw | CCAACCGAGGTGGATTTCGCGAAT | |
| | **MLY3015** | rv | CTAAGCTCTGACACCTCATCATTTCATC | |
| ***Meso3-1*** | **MLY3016** | fw | GGAGAGCAGGTGAAAGCAGTTGAG | |
| | **MLY3017** | rv | GATGTACATCCAATGGGAGATCTGTGACA | |
| ***Meso3-2*** | **MLY3018** | fw | CCGGCCATGATAAAGGAAAGATTGGAG | |
| | **MLY3019** | rv | GGTGCTTCAATCTTGACTATCTGGCC | |
| ***IAA14*** | **MLY2948** | fw | CCTCTCTTATTTTTTGTTATTACTGGAGTATATTATT | ISH |
| | **MLY2949** | rv | tgtaatacgactcactatagggcCTCTTTTCCCATTGACCTTATTATCAGC | |

**Table S9. List of primers used in this study.**

List of primers used for RT-qPCR (Figure S6) and *in situ* hybridization (ISH, Figure S6). The red nucleotides in primer MLY2949 indicate the T7 promoter for *in vitro* transcription.
